# Uptake of tumor-derived microparticles induces metabolic reprogramming of macrophages in the early metastatic lung

**DOI:** 10.1101/2022.11.15.516217

**Authors:** Kelly Kersten, Ran You, Sophia Liang, Kevin M. Tharp, Joshua Pollack, Valerie M. Weaver, Matthew F. Krummel, Mark B. Headley

## Abstract

The formation of a pre-metastatic niche is a critical step during the metastatic spread of cancer. One way by which primary tumors prime host cells at future metastatic sites is through local shedding of tumor-derived microparticles as a consequence of vascular sheer flow. However, it remains unclear how the uptake of such particles by resident immune cells affects their phenotype and function. Here we show that ingestion of tumor-derived microparticles by macrophages induces a rapid metabolic and phenotypic switch that is characterized by enhanced mitochondrial mass and function, increased oxidative phosphorylation and upregulation of cellular adhesion molecules resulting in reduced motility in the early metastatic lung. We show that this reprogramming event is dependent on signaling through the mTORC1, but not mTORC2 pathway, and is unique to uptake of tumor-derived microparticles. Together, these data support a mechanism by which uptake of tumor-derived microparticles induces reprogramming of macrophages to shape their fate and function in the early metastatic lung.

## Introduction

Metastatic spread is one of the main causes of cancer mortality. Yet, the mechanisms by which cancer cells disseminate from the primary tumor and successfully colonize distant organs remains largely unknown. Accumulating evidence demonstrates a pivotal role for immune cells in priming distant organs by creating a ‘pre-metastatic niche’ that supports the colonization and outgrowth of future incoming cancer cells. Partly, this is mediated by classical inflammatory monocytes and macrophages promoting metastatic spread of cancer through their regulation of tissue remodeling, angiogenesis and suppressing anti-tumor T cell responses^1–3^. Conversely, a population of non-classical patrolling monocytes prevented the formation of lung metastasis by engulfing tumor-derived material and promoting natural killer (NK) cell recruitment and activation^4^. The mechanisms by which tumor-derived factors instruct myeloid cells at these distant (pre-)metastatic sites remains undiscovered.

Monocytes and macrophages display a remarkable heterogeneity and plasticity that is highly dependent on spatial and temporal cues from their local microenvironment. Several studies have reported shedding of microparticles, including exosomes, as ways for tumors to instruct resident cells, including endothelial cells, fibroblasts and macrophages, at distant organs to create a niche that is then susceptible to disseminated cancer cells and supports metastatic growth^5–8^. We have previously reported that microparticles shed by primary B16F10 mouse melanomas are rapidly ingested by distinct waves of myeloid populations in the early metastatic lung, of which non-alveolar inflammatory macrophages were the most predominant^2^.

Here we examined the early consequences of ingestion of tumor-derived microparticles by macrophages, looking at their transcriptional profiles and changes in cellular metabolism to understand the nature of reprogramming of the myeloid compartment by early metastatic pioneers in the lung.

## Results

### Ingestion of tumor-derived microparticles induces transcriptional reprogramming of macrophages in the early metastatic lung

To study the dynamics of antigen-loading and how it affects the fate of myeloid populations in the early metastatic lung, we utilized an animal model for experimental metastasis by intravenous (i.v.) administration of a range of established cancer cell lines tagged with fluorescent marker ZsGreen (Fig. 1A, B). In line with previous results^2^, we found a diverse range of ZsGreen-antigen-loading among distinct myeloid populations in the lung 24 hours post i.v. injection of cancer cells (Fig. 1B and Supplemental Fig. 1A, B). Among them, non-alveolar inflammatory macrophages were the most predominant cell type ingesting ZsGreen^+^ material, accounting for ∼ 50% of all ZsGreen^+^ CD45^+^ immune cells at 24 hours post exposure (Fig. 1B and Supplemental Fig. 1B). This loading occurred similarly following injection of ZsGreen-tagged Lewis Lung Carcinoma (LLC), the poorly metastatic B16F1, as well as non-cancerous ZsGreen-tagged mouse embryonic fibroblasts (MEF) derived from *β-actin*^*Cre*^*;Ai6* mice (Fig. 1B). In contrast, following i.v. administration of fluorescent latex beads, patrolling monocytes represented the majority of bead-ingesting immune cells (Fig. 1B).

**Figure 1.**
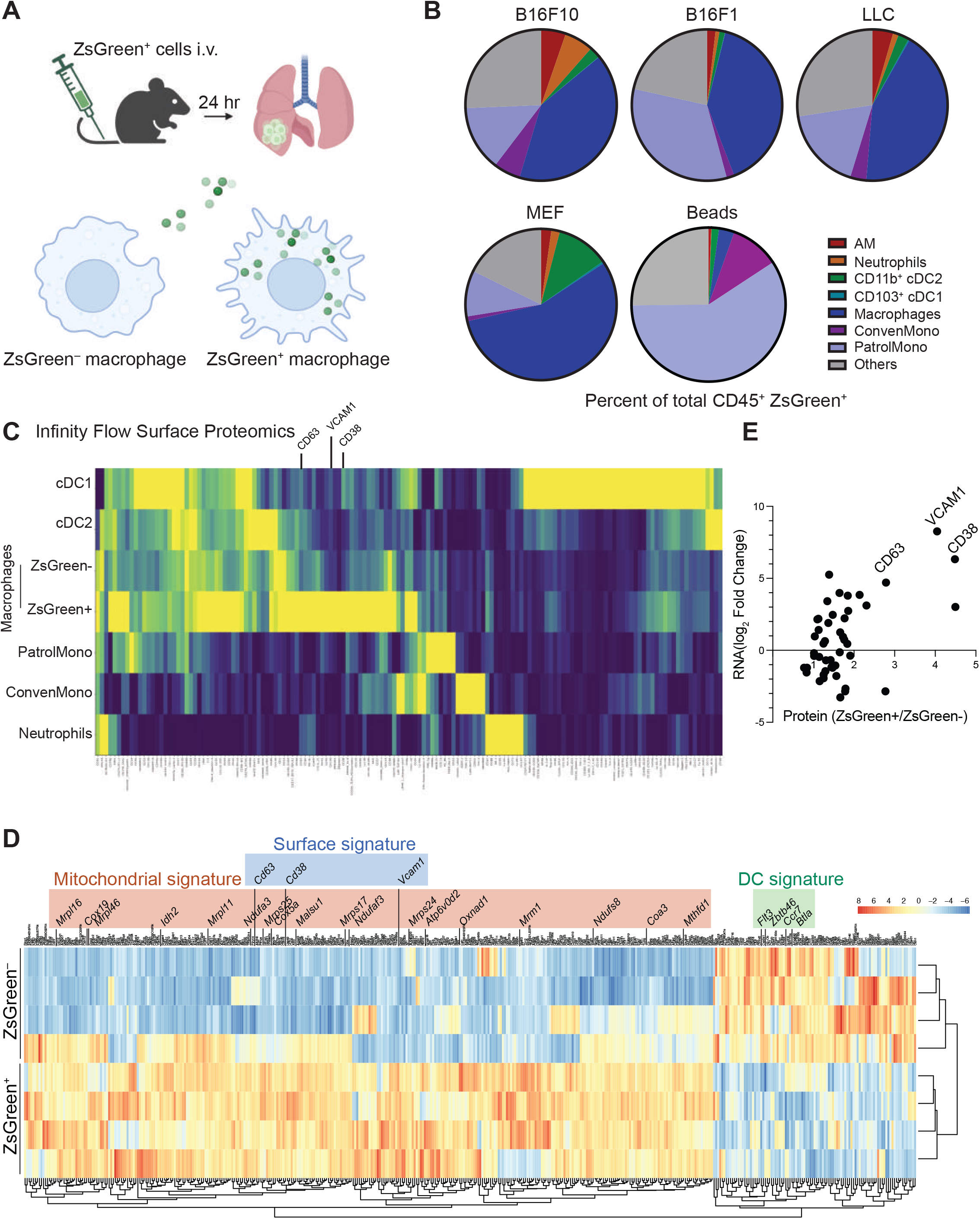
Uptake of tumor-derived microparticles induces transcriptional and phenotypic reprogramming of macrophages in the early metastatic lung. A) Schematic overview of experimental setup. Lungs were harvested 24 hours post intravenous administration of ZsGreen^+^ cells in the tail vein. B) Quantification of flow cytometric analysis of the uptake of ZsGreen^+^ microparticles by distinct CD45^+^ myeloid cell populations in the lung 24 hours post i.v. injection of cancer cells (B16F10, B16F1, or Lewis lung carcinoma (LLC)), mouse embryonic fibroblast (MEF) or FITC-labeled polystyrene beads. C) Heatmap displaying differential expression of surface protein markers on myeloid cell populations in the lung 24 hours post i.v. injection of B16ZsGreen cells using Infinity Flow Surface Proteomics. D) Heatmap displaying the top differentially expressed genes (DEG) in ZsGreen^+^ and ZsGreen^−^ macrophages isolated from lungs of mice 24 hours post i.v. injection of B16ZsGreen cells as described in panel A. DEG related to mitochondria (‘mitochondrial signature’), cellular adhesion markers (‘surface signature’) and dendritic cell functions (‘DC signature’) are color-coded separately. N = 4 per group. DEG genes were selected based on cut-off of FDR < 0.05. E) Correlation between proteomics (ratio of ZsGreen^+^/ZsGreen^−^) and RNA expression (log_2_ Fold Change) data in lung macrophages.

We next sought to ascertain whether gene and cell surface protein expression are modulated following ingestion of tumor-derived microparticles. We administered B16ZsGreen cells i.v. and 24 hours later performed cell surface proteomics on whole lung, using our recently developed Infinity Flow method^9^. We comprehensively examined the cell surface proteome of the subset of tumor-ingesting versus non-ingesting macrophages in direct comparison to other lung myeloid subsets (Fig. 1C and Supplemental Fig. 1C.) The tumor-ingesting macrophage population was identified based on high ZsGreen MFI (Supplemental Fig. 1D). This analysis revealed distinct phenotypic differences between macrophages and other lung myeloid cells, and further revealed an array of cell surface markers that differed in expression between ZsGreen^+^ and ZsGreen^−^ macrophages (Fig. 1C). We additionally utilized fluorescence activated cell sorting (FACS) to isolate lung macrophages that had ingested tumor-derived microparticles (ZsGreen^+^) versus those that had not (ZsGreen^−^) and performed RNA-sequencing to assess differential gene expression between the two macrophage populations. Analysis of differentially expressed genes (DEGs) revealed that uptake of microparticles induced a rapid transcriptional reprogramming characterized by the upregulation of a range of mitochondrial genes (referred to as ‘Mitochondrial signature’) as well as cellular adhesion markers (‘Surface signature’; e.g. VCAM1, CD38, CD63) (Fig. 1D). Importantly, this cell surface signature aligned with that revealed by Infinity Flow (Fig. 1E). Moreover, we found that expression of stimulatory dendritic cell (DC) genes^10,11^ – including *Flt3, Zbtb46, Ccr7* and *Btla* – was significantly downregulated in ZsGreen^+^ versus ZsGreen^−^ macrophages (‘DC signature’) (Fig. 1D). Together these data demonstrate that macrophages loaded with tumor-derived microparticles display distinct transcriptional and phenotypic programs as compared to non-loaded macrophages in the early metastatic lung.

### Phenotypic reprogramming of lung macrophages upon antigen-loading with tumor-derived microparticles

Based on the high concordance between protein and transcript levels, we selected CD63, VCAM1, and CD38 as a signature for further interrogation of the impact of tumor-ingestion on lung macrophage populations (Fig. 1E). We first validated the expression differences in independent flow cytometry experiments (Supplemental Fig. 2A.) Importantly, *in vitro* cultured bone marrow-derived macrophages (BMDM) similarly upregulated VCAM1, CD38, and CD63 in response to B16ZsGreen ingestion (Supplemental Fig. 2B,C), supporting the use of these proteins as a signature of ingestion-mediated modulation of macrophages. Antigen-loading (Supplemental Fig. 3A) and the resulting phenotypic change in lung macrophages after B16ZsGreen injection occurred similarly after administration of MEFZsGreen cells *in vivo*, however less pronounced (Fig. 2A, B). While VCAM1 and CD63 surface levels were only found augmented at 24 hours injection, CD38 expression levels were already substantially upregulated 2 hours after exposure (Supplemental Fig. 3C). Uptake of FITC-labeled beads modestly increased the expression of VCAM1 and CD63 on lung macrophages, but the levels were significantly lower following ingestion of B16-derived microparticles (Supplemental Fig. 4A-H).

**Figure 2.**
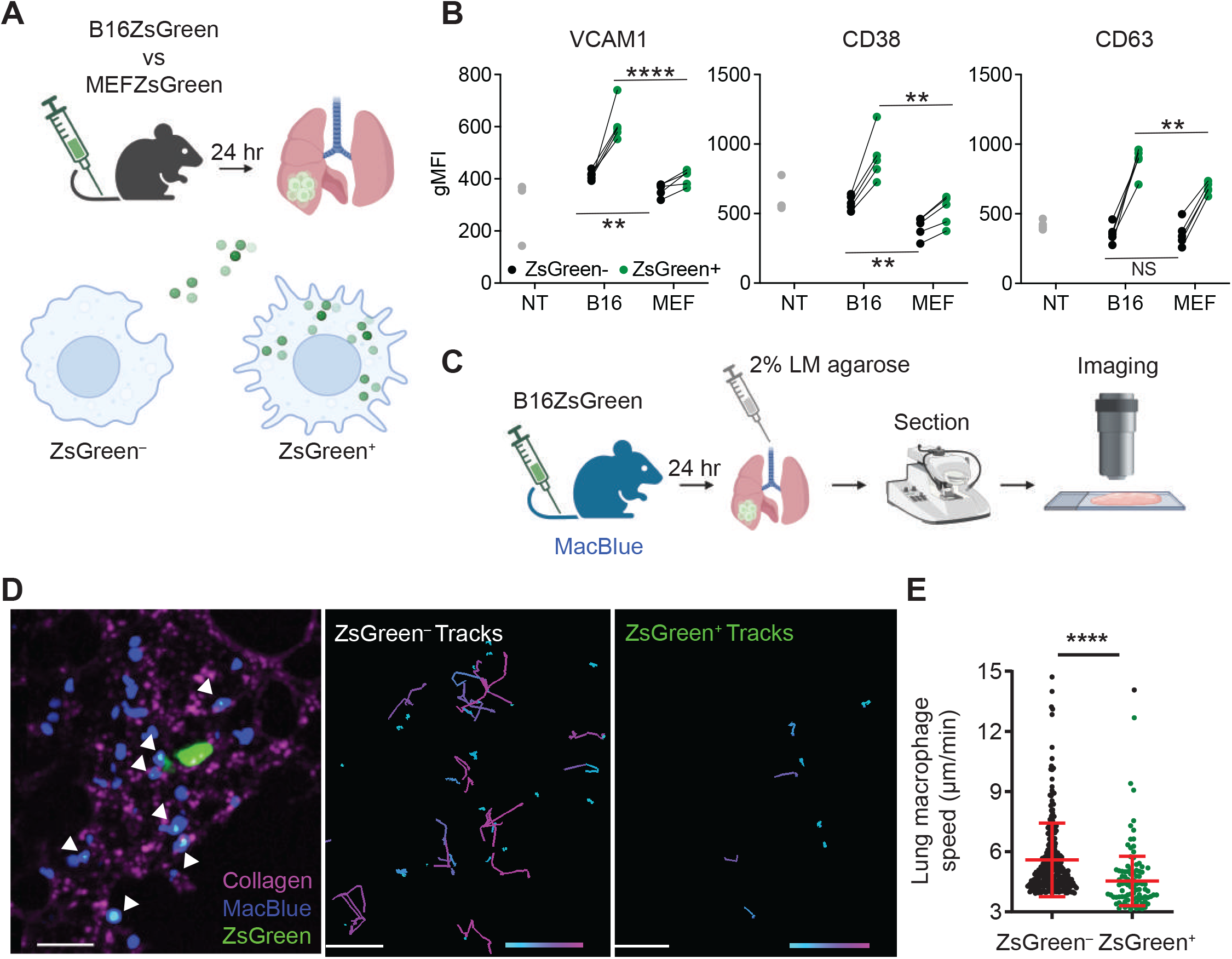
Ingestion of tumor-derived microparticles induces phenotypic reprogramming of macrophages in the early metastatic lung. A) Schematic overview of experimental setup. Lungs were harvested 24 hours post intravenous administration of B16ZsGreen^+^ cells or MEFZsGreen^+^ cells in the tail vein. B) Quantification of VCAM1, CD38 and CD63 in ZsGreen^-^ and ZsGreen^+^ lung macrophages 24 hours post i.v. injection of B16ZsGreen (B16), MEFZsGreen (MEF) by flow cytometry. C) Experimental layout of live precision-cut tissue slice imaging of lungs of MacBlue mice harvested post i.v. injection of B16ZsGreen. D) Representative image and track displacement display of ZsGreen^+^ versus ZsGreen^-^ eCFP^+^ cells over 1 hour in live lung slices from MacBlue mice at 24 hours post i.v. injection of B16ZsGreen cells. White arrows indicate ZsGreen-microparticle-ingested-eCFP^+^ cells. Track length is ranged from 3μm (cyan) to 35μm (pink). E) Quantification of the instant speed of eCFP^+^ cells in live lung slices. Data are pooled from three different ROIs and are representative of two independent experiments. **** p < 0.0001, ** p < 0.01 as determined by Student’s t-test.

To assess whether the upregulation of adhesion molecules in ZsGreen^+^ versus ZsGreen^−^ macrophages affected macrophage phenotype in the lung, we performed two-photon imaging in live lung slices obtained from MacBlue mice – in which eCFP expression is driven by a modified *c-Fms* promoter allowing the visualization of monocytes, monocyte-derived macrophages and a small proportion of neutrophils^12^. 24 hours after i.v. injection of B16ZsGreen cells (Fig. 2C), ZsGreen^+^ eCFP^+^ cells exhibited substantially lower motility compared to ZsGreen^−^ eCFP^+^ cells in the lung (Fig. 2D, E), an effect that might be due to reduced cell-intrinsic motility, a change of location relative to tissue, or upregulated expression of adhesion markers.

### Tumor-derived antigen-loading drives mitochondrial reprogramming in lung macrophages

To further study macrophage phenotypic reprogramming in the early metastatic lung, we performed gene set enrichment analysis (GSEA) on our RNA-seq dataset (Fig. 1D). This revealed four major pathway networks closely associated with translation, protein processing, catabolic processes and mitochondrial respiration, that were highly enriched in ZsGreen^+^ macrophages as compared to ZsGreen^−^ macrophages (Fig. 3A and Table S1).

**Figure 3.**
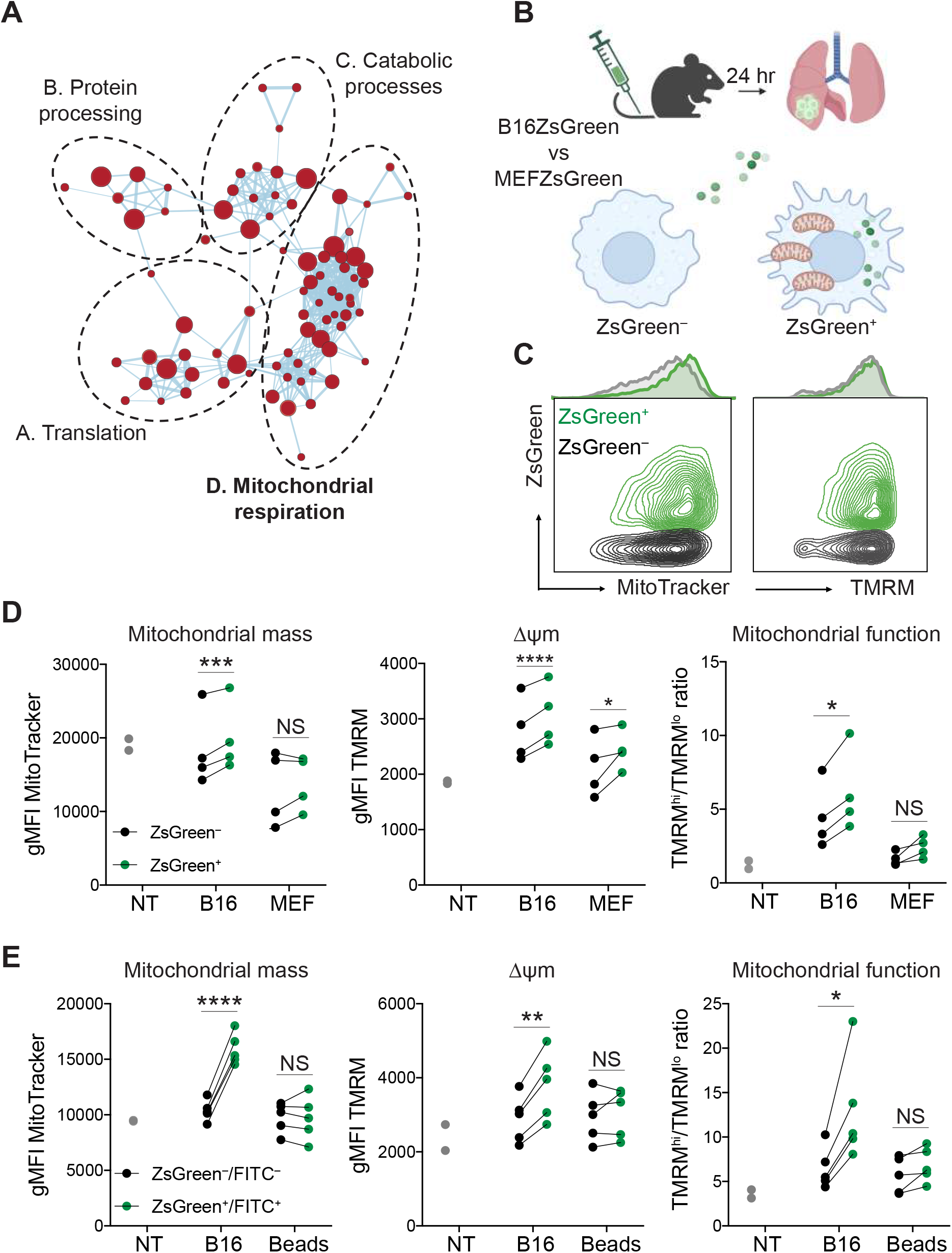
Tumor-derived antigen-loading drives mitochondrial reprogramming in lung macrophages. A) Cytoscape network analysis of enriched pathways in ZsGreen^+^ versus ZsGreen^-^ lung macrophages. Also see Supplemental Table 1 for a list of the identity of pathways in each cluster. B) Schematic overview of experimental setup. Lungs were harvested 24 hours post i.v. injection of B16ZsGreen or MEFZsGreen in the tail vein. C) Representative contour plots and histograms of MitoTracker DeepRed and TMRM staining of ZsGreen^+^ and ZsGreen^-^ lung macrophages. D-E) Quantification of mitochondrial mass (MitoTracker DeepRed), mitochondrial membrane potential (TMRM), and mitochondrial function (ratio of the percentage of TMRM^hi^ to TMRM^lo^ cells) in ZsGreen^-^ and ZsGreen^+^ lung macrophages 24 hours post i.v. injection of B16ZsGreen (B16), MEFZsGreen (MEF), FITC-labeled polystyrene beads (Beads) or non-tumor challenged (NT) mice. Data are representative of two or three independent experiments; **** p < 0.0001, *** p < 0.001, ** p < 0.01, * p < 0.05 as determined by the Paired Student’s *t*-test.

Following up on the ‘Mitochondrial signature’ induced in antigen-loaded macrophages (Fig. 1D), we quantified the variation in mitochondrial mass (indicated by MitoTracker DeepRed staining) and mitochondrial membrane potential (ΔΨm; determined by fluorescence of the potential-sensitive tetramethylrhodamine methyl (TMRM) dye) after *in vivo* antigen-loading upon i.v. injection with B16ZsGreen or MEFZsGreen cells (Fig. 3B). ZsGreen^+^ macrophages exhibited a significant increase in mitochondrial mass, ΔΨm and TMRM^hi^/TMRM^low^ ratio when compared to ZsGreen^−^ macrophages from the same mice, consistent with our GSEA analysis (Fig. 3C, D). Notably, this was not the case for macrophages loaded with MEF-derived microparticles or FITC-labeled beads (Fig. 3D, E), suggesting that the mitochondrial reprogramming of lung macrophages is induced exclusively upon antigen-loading with tumor-derived material.

To extend this analysis to oxygen consumption rate (OCR), we developed an *in vitro* surrogate assay in which bone marrow-derived macrophages (BMDM) are cultured with B16ZsGreen tumor cells, and ZsGreen^+^ and ZsGreen^−^ BMDM are isolated for study after 24 hours (Fig. 4A). Consistent with the increased mitochondrial density and membrane potential observed in macrophages directly *ex vivo* (Fig. 3C-E), ZsGreen^+^ BMDM from this *in vitro* exposure showed a significantly higher OCR with higher basal and maximal respiration, as well as increased ATP production when compared to ZsGreen^−^ BMDM or non-tumor (NT) BMDM, which had never encountered tumor cells (Fig. 4B, C). Moreover, the spare respiratory capacity (SRC) was increased in ZsGreen^+^ BMDM, suggesting an improved capability to respond to sudden changes in energy demand (Fig. 4C). To determine whether this was phenocopied in cells that were programmed through exposure *in vivo*, we isolated lung macrophages from B16ZsGreen-challenged mice and compared their OCR to lung macrophages from unchallenged (NT) mice. In line with our *in vitro* data, lung macrophages that encountered tumor material showed increased OCR compared to NT mice (Supplemental Fig. 5A, B). To test whether this metabolic shift in lung macrophages is dictated by the source of antigen, we sorted ZsGreen^+^ and ZsGreen^−^ macrophages from lungs of either B16ZsGreen or MEFZsGreen-injected mice (Fig. 4D) and determined their ATP production as a direct measurement of mitochondrial function. While antigen-loading of macrophages with B16-derived microparticles induced a significant increase in ATP production in ZsGreen^+^ versus ZsGreen^−^ macrophages, loading with MEF-derived microparticles did not (Fig. 4E). Taken together, our data show that antigen-loading with tumor-derived material specifically enhanced mitochondrial respiration and increased ATP production in lung macrophages.

**Figure 4.**
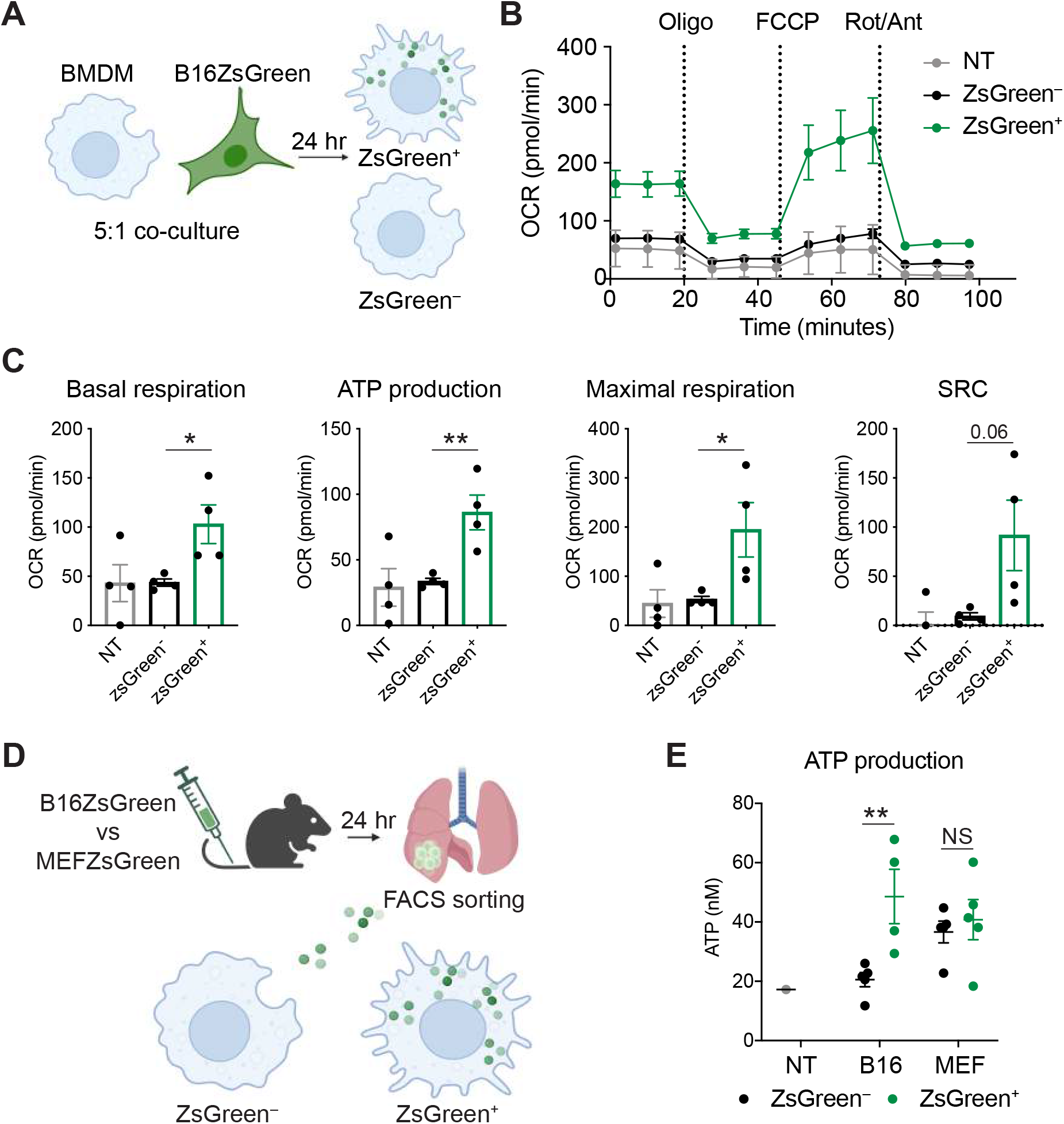
Metabolic reprogramming of lung macrophages upon ingestion of tumor-derived microparticles is characterized by enhanced oxidative phosphorylation. A) Schematic overview of experimental layout. Bone marrow-derived macrophages (BMDM) and B16ZsGreen cells were co-cultured in 5:1 ratio for 24 hours after which CD11b^+^ ZsGreen^-^, ZsGreen^+^ or non-tumor challenged (NT) BMDM were isolated using FACS-sorting and used for downstream analysis. B) FACS-sorted CD11b^+^ ZsGreen^-^, ZsGreen^+^ and NT BMDM were plated into Seahorse XFe24 plates and oxygen consumption rate (OCR) was determined using Seahorse extracellular flux assay with Oligomycin (Oligo), FCCP and rotenone and antimycin A (Rot/Ant) added sequentially. C) Quantification of basal respiration, ATP production, maximal respiration, and spare respiratory capacity (SRC) from extracellular flux assay. D) Schematic overview of experimental layout of isolation of ZsGreen^-^ and ZsGreen^+^ macrophages from lungs harvested 24 hours post i.v. injection of B16ZsGreen or MEFZsGreen cells by FACS-sorting. E) Quantification of ATP production in FACS-sorted ZsGreen^-^ and ZsGreen^+^ macrophages described in D determined by luminescence detection per manufacturer instructions. Data are representative of two independent experiments; all data represented as mean ± SEM; ** p < 0.01, * p < 0.05 as determined by the Unpaired Student’s *t*-test (C) or two-way (E) ANOVA and Sidak’s multiple comparison test.

### Macrophage reprogramming upon ingestion of tumor-derived microparticles is mTORC-dependent

To study the pathways regulating reprogramming of lung macrophages after ingestion of tumor-derived microparticles, we performed Hallmark Pathway Analysis comparing the transcriptional profile of ZsGreen^−^ and ZsGreen^+^ macrophages. We identified several metabolism-associated pathways among which “Oxidative phosphorylation” was highly enriched in ZsGreen^+^ versus ZsGreen^−^ macrophages (Fig. 5A). The mammalian target of rapamycin (mTOR) – the catalytic subunit of the two distinct complexes mTORC1 and mTORC2 – is a central regulator of cellular metabolism and has been shown to drive mitochondrial biogenesis and oxidative function^13,14^. Macrophage activation is dependent on mTOR and roles for both mTORC1^15^ and mTORC2^16,17^ have been implicated. In our data, mTORC1 (mammalian target of rapamycin complex 1) signaling was among the enriched pathways in ZsGreen^+^ versus ZsGreen^−^ macrophages (Fig. 5A). In line with this, detailed analysis of the genes that constitute the mTORC1 pathway revealed upregulation of genes encoding phospholipase D1 and 3 (*Pld1* and *Pld3*) and protein synthesis genes *Eif4ebp1, Rps23* and *Ptpa* (Fig. 5B). In contrast, expression of positive regulator gene of AMP-activated protein kinase (AMPK), *Prkab2*, was downregulated in ZsGreen^+^ versus ZsGreen^−^ macrophages (Fig. 5B), indicating that the AMPK pathway, as a negative upstream regulator of mTOR signaling^18^, was repressed. In line with our transcriptional data, we detected significantly higher levels of phosphorylated (p)4EBP1 and pS6 protein in ZsGreen^+^ versus ZsGreen^−^ macrophages by flow cytometry, confirming the activation of mTORC1 signaling upon ingestion of tumor-derived microparticles (Fig. 5C).

**Figure 5.**
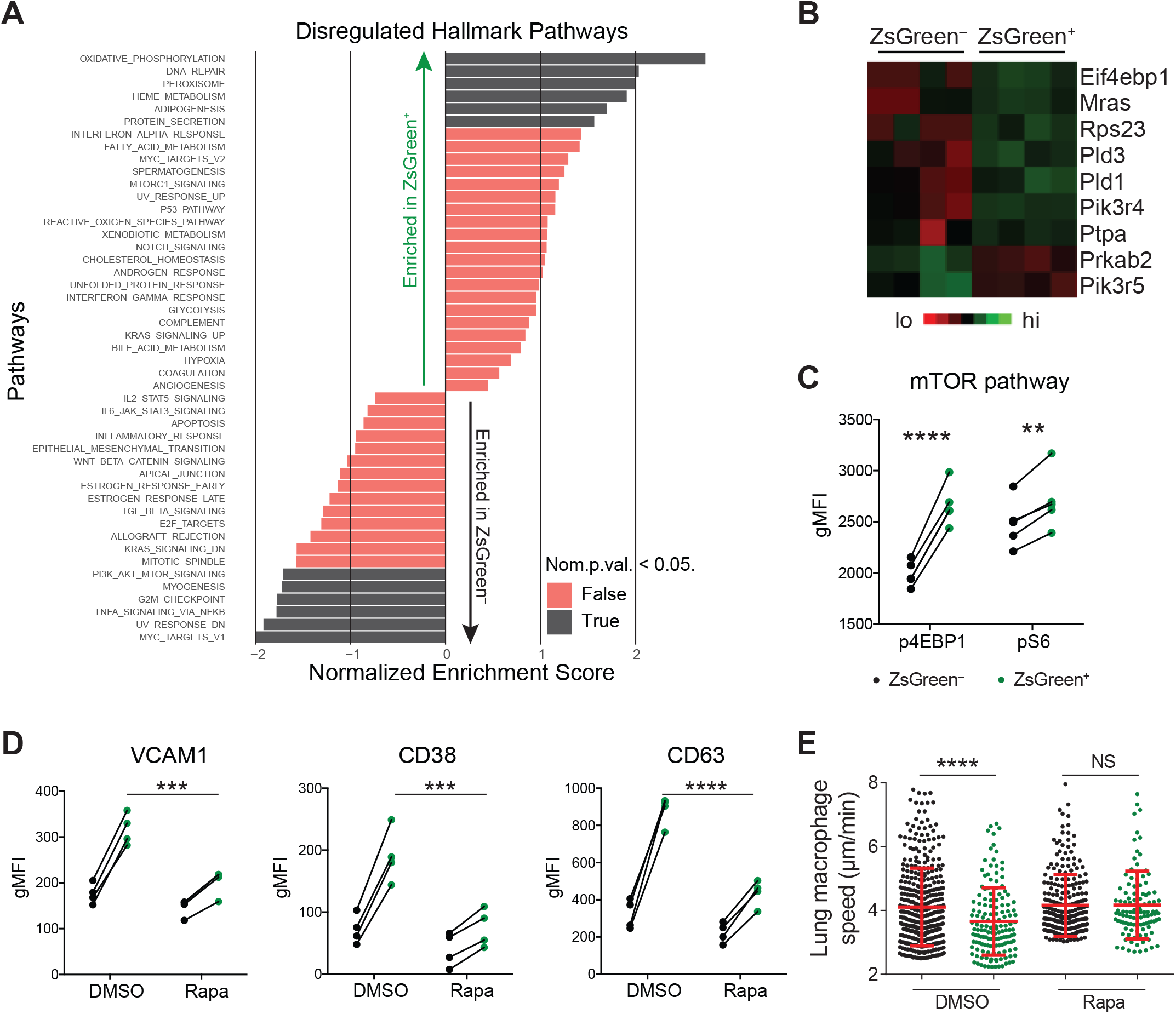
Phenotypic reprogramming of tumor microparticle-ingesting macrophages is mTORC1-dependent. A) Hallmark Pathway Analysis showing pathways upregulated (Normalized Enrichment Score >0) or downregulated (Normalized Enrichment Score <0) in ZsGreen^+^ versus ZsGreen^-^ macrophages. B) Heatmap displaying expression of single DEGs in mTORC1 pathway in ZsGreen^-^ versus ZsGreen^+^ macrophages. C) Quantification of phospho-(p)4EBP1 and pS6 expression on ZsGreen^-^ versus ZsGreen^+^ macrophages 24 hours post i.v. injection of B16ZsGreen by intracellular flow cytometry. D) Quantification of VCAM1, CD38 and CD63 on ZsGreen^-^ versus ZsGreen^+^ lung macrophages harvested 24 hours post i.v. injection of B16ZsGreen. Intranasal DMSO or rapamycin (0.4mg/kg) was administered twice, once on the day before and once on the day of tumor injection. E) Quantification of the instant speed of ZsGreen^-^ versus ZsGreen^+^ eCFP^+^ cells in live lung slices from the same mice depicted in D. Data are pooled from three different ROIs for each group. Data are representative of two independent experiments; **** p < 0.0001, *** p < 0.001 as determined by Paired Student’s *t*-test (C), two-way ANOVA and Sidak’s multiple comparison test (D), and Unpaired Student’s *t*-test (E).

To study whether the phenotypic switch in lung macrophages in response to ingestion of tumor-derived microparticles is regulated by mTOR, we treated mice intranasally with rapamycin to inhibit mTOR activation locally in the lungs of B16ZsGreen-injected mice. We found that upregulation of VCAM1, CD38 and CD63 expression on ZsGreen^+^ lung macrophages was significantly abolished upon rapamycin treatment as compared to DMSO-treated mice (Fig. 5D). Through imaging, we also found that ZsGreen^+^ myeloid cells in rapamycin-treated mice failed to arrest on lung parenchyma in contrast to DMSO-treated controls (Fig. 5E), suggesting that macrophage phenotypic reprogramming upon the ingestion of tumor-derived microparticles is dependent on mTORC signaling.

### mTORC1 is required for metabolic and phenotypic reprogramming of macrophages in the early metastatic lung

Because intranasal administration of rapamycin does not exclusively target macrophages, we undertook a genetic approach to test the various contributions of the two mTORC signaling complexes; genetic loss of Regulatory-associated protein of mTOR (Raptor) results in loss of the mTORC1 protein complex, while the mTORC2 complex is lost in mice lacking Rapamycin-insensitive companion of mTOR (Rictor)^18^. Using mice bearing conditional alleles *Raptor*^*f/f*^ and *Rictor*^*f/f*^ together with a Cre-recombinase under the control of the lysozyme 2 gene, we generated mice with myeloid-specific deletion of mTORC1 (*Rptr*^*f/f*^*LysM*^*Cre*^) and mTORC2 (*Rctr*^*f/f*^*LysM*^*Cre*^) complexes, respectively (Fig. 6A). While deficiency of mTORC1 or mTORC2 did not affect the proportion of lung macrophages nor their ability to ingest ZsGreen^+^ tumor-derived microparticles (Supplemental Fig. 6A, B), we detected a significant reduction in the ability to increase mitochondrial mass, membrane potential and mitochondrial function in ZsGreen^+^ versus ZsGreen^-^ macrophages in *Rptr*^*f/f*^*LysM*^*Cre*^ mice, as compared to WT littermates or *Rctr*^*f/f*^*LysM*^*Cre*^ mice (Fig. 6B), demonstrating that mTORC1 signaling specifically, is required for the metabolic switch in macrophages upon antigen-loading.

**Figure 6.**
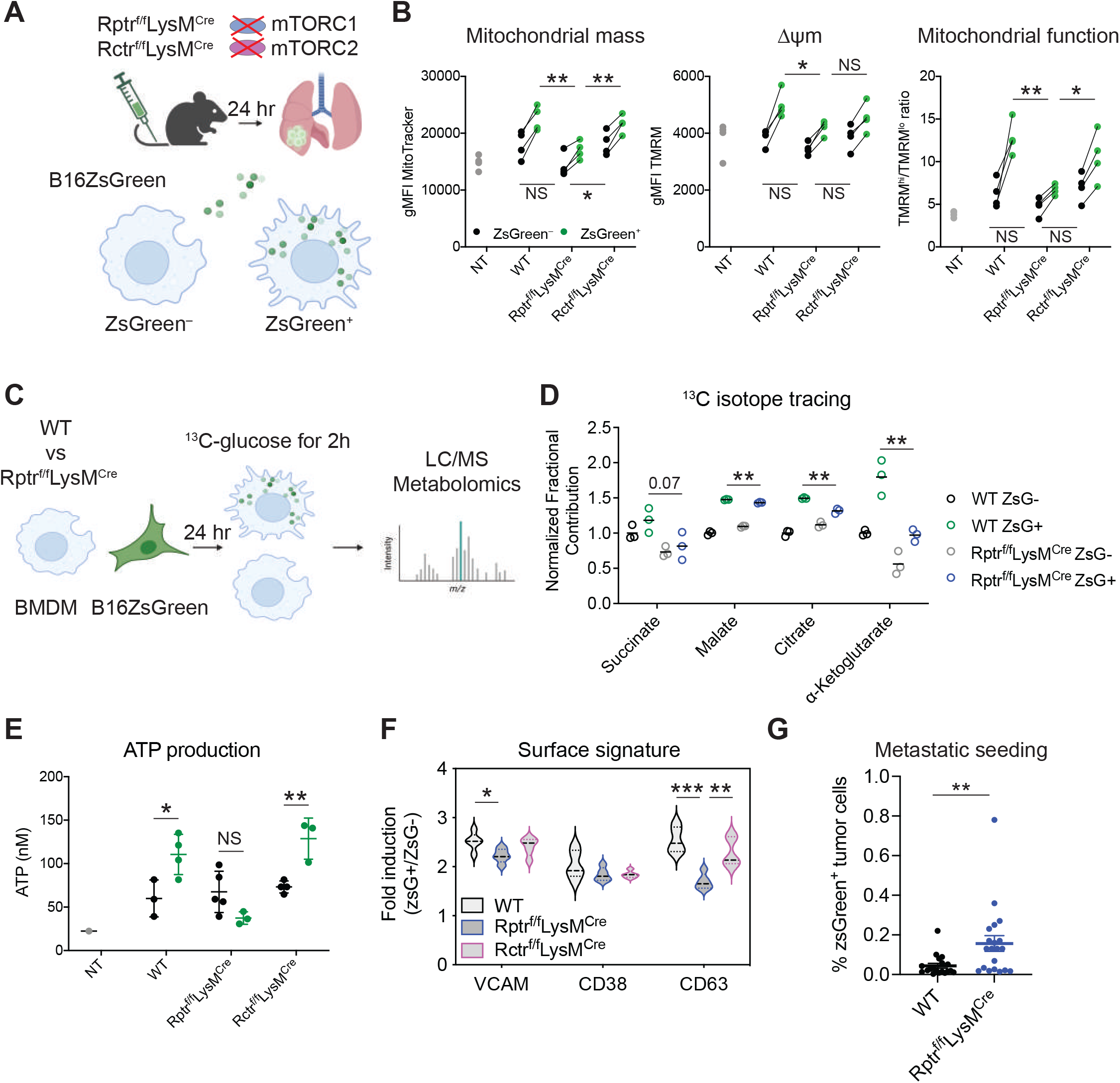
mTORC1 is required for metabolic and phenotypic reprogramming of macrophages in the early metastatic lung. A) Experimental layout of B16ZsGreen administration in mTORC1-deficient *Rptr*^*f/f*^*/LysM*^*Cre*^, mTORC2-deficient *Rctr*^*f/f*^*/LysM*^*Cre*^ mice and wild type (WT) littermates. B) Quantification of mitochondrial mass (MitoTracker DeepRed), mitochondrial membrane potential (TMRM), and mitochondrial function (ratio of the percentage of TMRM^hi^ to TMRM^lo^ cells) in ZsGreen^-^ and ZsGreen^+^ lung macrophages 24 hours post i.v. injection of B16ZsGreen in indicated groups of mice by flow cytometry. Statistical differences were determined by Student’s Unpaired t-test. C) Schematic overview of metabolomics experiment. WT or *Rptr*^*f/f*^*/LysM*^*Cre*^ BMDM were co-cultured with B16ZsGreen at a 5:1 ratio for 24 hours followed by FACS-sorting of ZsGreen^-^, ZsGreen^+^ and non-tumor challenged (NT) BMDM. Cells were rested for 1 hour and media was substituted with media containing isotopically labeled glucose (^13^C) for 2 hours ^19^ upon which cells were fixed and metabolites were extracted for LC/MS analysis. D) Normalized fractional contribution of incorporation of ^13^C-glucose in selected TCA cycle metabolites in FACS-sorted ZsGreen^-^, ZsGreen^+^ and NT wild type (WT) or mTORC1-deficient *Rptr*^*f/f*^*/LysM*^*Cre*^ BMDM. E) Quantification of ATP production in ZsGreen^-^ or ZsGreen^+^ macrophages isolated from lungs of indicated groups of mice. F) Fold increase of VCAM1, CD38 and CD63 expression on ZsGreen^+^ over ZsGreen^-^ lung macrophages in indicated groups of mice. N = 3-5 per group. G) Quantification of CD45^−^ ZsGreen^+^ tumor cells of total live cells in lungs 24 hours post i.v. injection of B16ZsGreen indicative of metastatic seeding by flow cytometry (n=19-20 mice/group; pooled data from 5 independent experiments). All data are representative of at least two independent experiments; ** p < 0.01, * p < 0.05 as determined by Unpaired Student’s *t*-test (B, D, F), two-way ANOVA and Sidak’s multiple comparison test (E), Mann-Whitney U-test (G).

To directly test how mitochondrial metabolism in macrophages was affected after uptake of tumor-derived microparticles, we sorted ZsGreen^+^ and ZsGreen^−^ BMDM after 24 hours of co-culture with B16ZsGreen cells and monitored their metabolism of isotopically labeled glucose (^13^C) for 2 hours^19^ (Fig. 6C and Supplemental Fig. 6C). Ingestion of ZsGreen^+^ microparticles by wild type BMDM (WT ZsGreen^+^ vs WT ZsGreen^—^) increased mitochondrial metabolism of glucose indicated by increased incorporation of ^13^C into TCA cycle intermediates (Supplemental Fig. 6C), which corroborates our cellular respirometry data obtained with our Seahorse assays (Fig. 4B, C and Supplemental Fig. 5A, B). This process was attenuated in mTORC1-deficient BMDM derived from *Rptr*^*f/f*^*LysM*^*Cre*^ mice (Supplemental Fig. 6C). Specifically, we found reduced isotopic labeling of succinate, malate, citrate and alpha-ketoglutarate in ZsGreen^+^ *Rptr*^*f/f*^*LysM*^*Cre*^ BMDM as compared to ZsGreen^+^ WT BMDM (Fig. 6D).

To validate our findings from the isotopic tracing experiment *in vivo*, we sorted ZsGreen^+^ and ZsGreen^−^ lung macrophages from WT, *Rptr*^*f/f*^*LysM*^*Cre*^ and *Rctr*^*f/f*^*LysM*^*Cre*^ mice 24 hours after i.v. injection of B16ZsGreen cells (Fig. 6A), and directly measured their ATP production. Consistently, the increased ATP production in ZsGreen^+^ versus ZsGreen^-^ macrophages as observed in WT mice, was completely abolished in *Rptr*^*f/f*^*LysM*^*Cre*^ mice, while the absence of mTORC2 in macrophages from *Rctr*^*f/f*^*LysM*^*Cre*^ mice displayed ATP production similar to controls (Fig. 6E). Moreover, upregulation of VCAM1 and CD63, but not CD38, on ZsGreen^+^ macrophages was significantly lower in *Rptr*^*f/f*^*LysM*^*Cre*^ mice when compared to WT and *Rctr*^*f/f*^*LysM*^*Cre*^ mice (Fig. 6F). We thus conclude that metabolic and phenotypic reprogramming of lung macrophages after uptake of tumor material is dependent on mTORC1 signaling.

Finally, to test the functional significance of mTORC1-dependent macrophage reprogramming for the clinical outcome of metastatic disease, we i.v. injected *Rptr*^*f/f*^*LysM*^*Cre*^ mice and WT littermates with B16ZsGreen and allowed lung metastasis to develop. Interestingly, at 24 hours post i.v. injection with B16ZsGreen we found a modest but significant increase in the proportion of ZsGreen^+^ CD45^−^ cells in the lungs of *Rptr*^*f/f*^*LysM*^*Cre*^ mice as compared to WT littermates (Fig. 6F), suggestive of increased metastatic seeding. However, by 2-3 weeks of lung colonization, the total number of metastatic nodules (Supplemental Fig. 6D) or the metastatic burden (measured as the area of the lung affected by metastasis) (Supplemental Fig. 6E) were not affected by macrophage-specific mTORC1-deficiency. This normalization may be a result of local changes in nutrient availability as a result of rapidly growing tumor lesions dominating the lung metastatic microenvironment.

## Discussion

A number of studies have implicated the important role of macrophages in the metastatic spread of cancer^20,21^. However, it remains unclear how the early interactions between disseminated cancer cells and macrophages in pre-metastatic tissues dictate successful colonization of secondary organs. Previously, we and others have shown that tumors actively shed microparticles (including exosomes) into the circulation that migrate to and condition pre-metastatic sites such as the lungs^2,6–8,22^. In this work, we have demonstrated that ingestion of these tumor-derived microparticles induces a metabolic and phenotypic switch in non-resident macrophages in the early metastatic lung, that is highly dependent on mTORC1 signaling. We found that this reprogramming event is specific for the uptake of tumor-derived material. Thus, these initial changes might be an important step into shaping macrophage function during the metastatic spread of cancer.

A growing number of studies have highlighted the importance of metabolism in dictating macrophage activation and function^23–27^. However, a large number of contradictory observations have complicated their interpretation, possibly caused by fundamental differences between *in vitro* and *in vivo* experimental settings and the fact that the binary anti-tumor or ‘M1-like’ versus pro-tumor ‘M2-like’ polarization model does not accurately reflect the heterogeneity of macrophages, especially in the context of cancer^28–30^. For example, mTORC1-driven differentiation of monocytes into M2-like TAM was found to promote angiogenesis and tumor growth in a liver cancer model^31^. In contrast, Wenes *et al*. reported that activation of mTORC1 – through deletion of REDD1 – in hypoxic TAM enhanced a switch to glycolysis, normalized the tumor vasculature restoring oxygenation and prevented formation of metastasis^32^. Our data add another layer that might explain some of these seemingly conflicting results. Our work shows that the initial ingestion of tumor-derived material induces rapid mTORC1 activation in macrophages driving metabolic programs characterized by enhanced oxidative phosphorylation that support an anti-metastatic function in the pre-metastatic lung (Fig. 6G). However, this anti-tumor phenotype seems to be lost upon prolonged exposure to growing metastatic lesions (Supplemental Fig. 6D, E). It is plausible that the high metabolic demand of tumor cells in overt metastatic lungs enforces temporal changes in the local composition of oxygen and nutrients that result in spatiotemporal metabolic changes driving a shift in macrophage function from an anti-tumor to a pro-tumor state. Adding to the complexity, the metabolic changes in immune cells observed in tumor studies seem to be highly dependent on context, including the timing of experimental analysis and the type of preclinical cancer model used. Thus, further studies are needed to understand how macrophage function is shaped by changes in metabolic states over time in growing tumors.

Manipulation of macrophages has been an interesting avenue for therapeutic intervention to prevent or slow down tumor progression and metastatic spread in a substantial proportion of cancer patients. However, the success of myeloid cell-targeting compounds in clinical trials has been limited^33^. A major caveat in developing macrophage-targeting compounds is most likely a lack of specificity caused by the extremely heterogeneous and plastic nature of these cells. Ideally, one would target pro-tumor macrophage states and leave the anti-tumor macrophages unaffected. Based on our study and others’, it would be interesting to identify ways of targeting certain metabolic states in macrophages to selectively suppress the pro-tumor function of macrophages and enhance their anti-tumor function to work in conjunction with other anti-cancer therapies.

## Supporting information

Key Resources

## Acknowledgements

We thank all members of the Krummel laboratory for discussion and support. We also thank the ImmunoX CoLabs at UCSF for technical assistance and support. We thank Johanna ten Hoeve and the UCLA Metabolics Center for technical assistance and performing metabolic tracing experiments. Figures were created using BioRender.com. Funding: K.K. was supported by a Rubicon postdoctoral fellowship 019.163LW.006 from the Netherlands Organization for Scientific Research (NWO), and a Parker Scholar Award from the Parker Institute for Cancer Immunotherapy. We acknowledge the UCSF Parnassus Flow Core (RRID:SCR_018206 and DRC Center Grant NIH P30 DK063720) for assistance and using their instruments and services. This work was supported by NIH/NCI U54CA163123, 5U01CA217864, R21CA191428 and R01CA197363 to M.F.K.

## Author contributions

K.K. and R.Y. designed and conducted most experiments, data analysis, and drafted the manuscript. S.L. and M.B.H. generated ZsGreen-expressing cell lines, performed RNA-seq and Infinity Flow experiments. J.P. analyzed RNA-seq data. K.M.T. assisted and analyzed the metabolic isotope tracing experiment under supervision of V.M.W.. M.F.K. designed the experiments, interpreted data, and with other co-authors, developed the manuscript.

## Supplemental Figure Legends

**Supplemental Figure 1.**
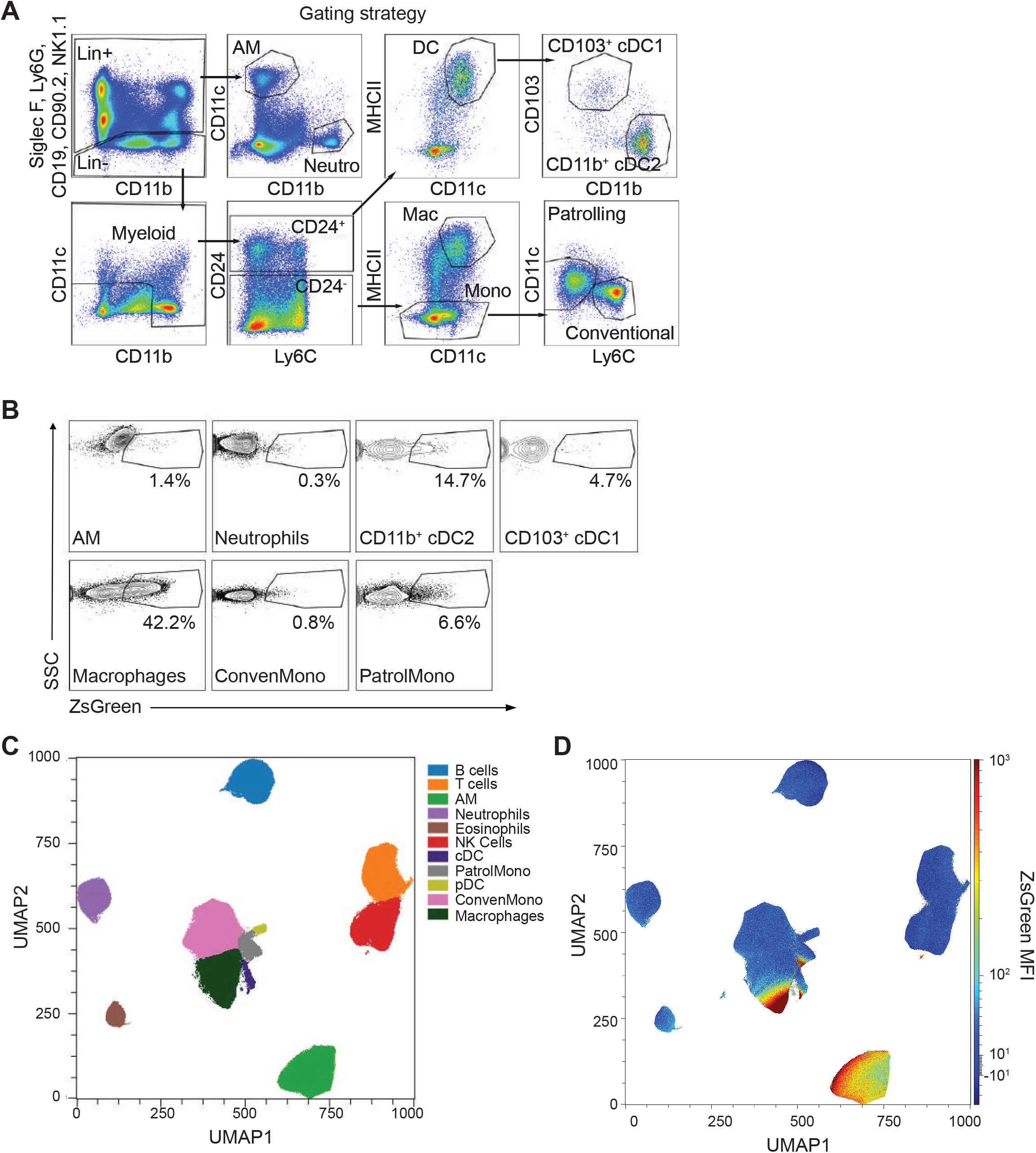
Uptake of tumor-derived microparticles by distinct populations of myeloid cells in the lung. A) Representative dot plots depicting the gating strategy for distinct myeloid populations in the lung. B) Representative contour plots showing the proportion of ZsGreen-uptake by the indicated myeloid populations in the lung. C) UMAP displaying distinct immune cell populations in the lung 24 hours after injection of B16ZsGreen i.v. using Infinity Flow Surface Proteomics. D) UMAP overlayed with ZsGreen Mean Fluorescence Intensity (MFI) by Infinity Flow Surface Proteomics.

**Supplemental Figure 2.**
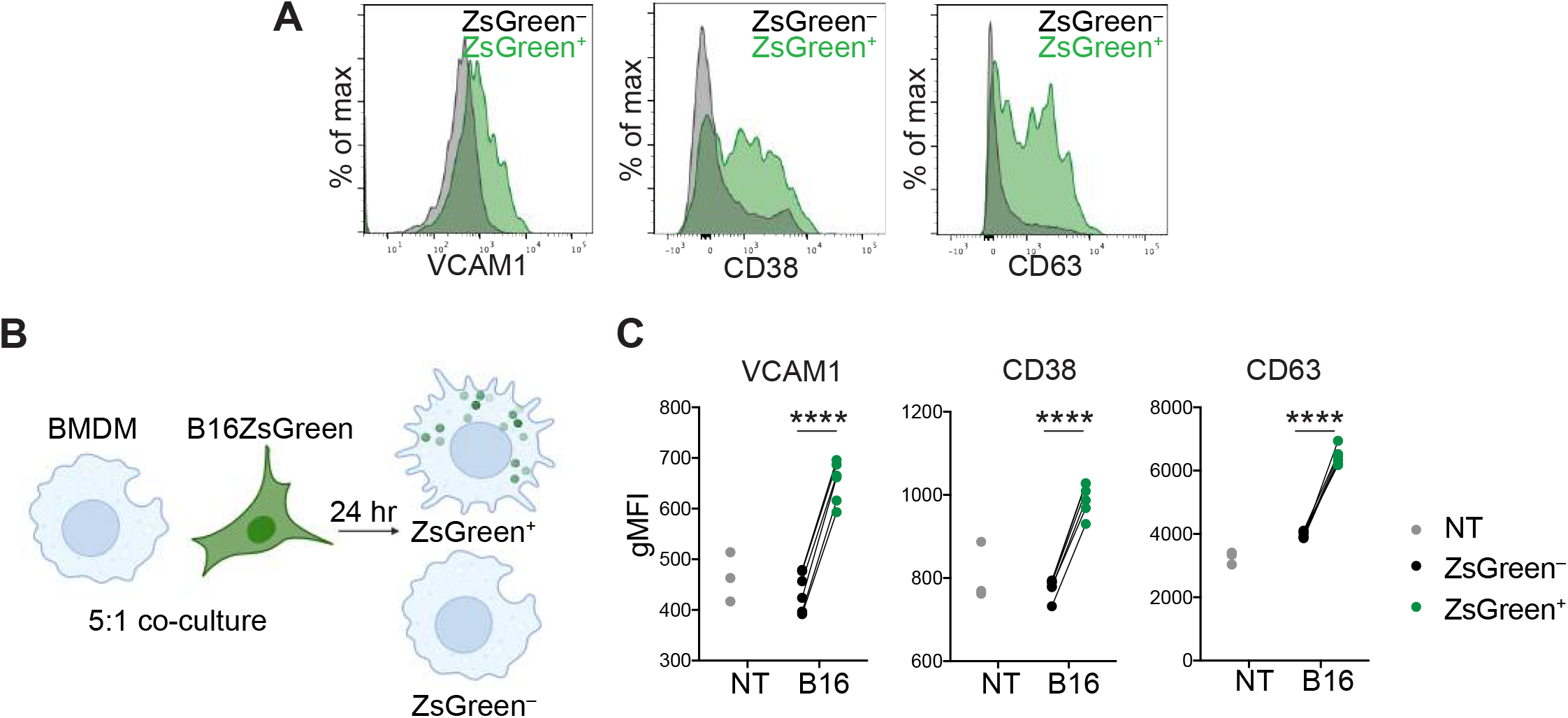
Ingestion of tumor-derived microparticles induces a phenotypic change in macrophages. A) Representative histogram of expression of VCAM1, CD38 and CD63 on ZsGreen^+^ versus ZsGreen^-^ lung macrophages *in vivo* 24 hours after injection of B16ZsGreen i.v. by flow cytometry. B) Experimental layout of *in vitro* coculture system of bone marrow-derived macrophages (BMDM) and B16ZsGreen cells at a 5:1 ratio for 24 hours prior to flow cytometric analysis of BMDM phenotype. C) Quantification of VCAM1, CD38 and CD63 expression on ZsGreen^-^, ZsGreen^+^ or non-tumor challenged (NT) BMDM as determined by flow cytometry. **** p < 0.0001 as determined by Paired Student’s *t*-test. Data are representative of three independent experiments.

**Supplemental Figure 3.**
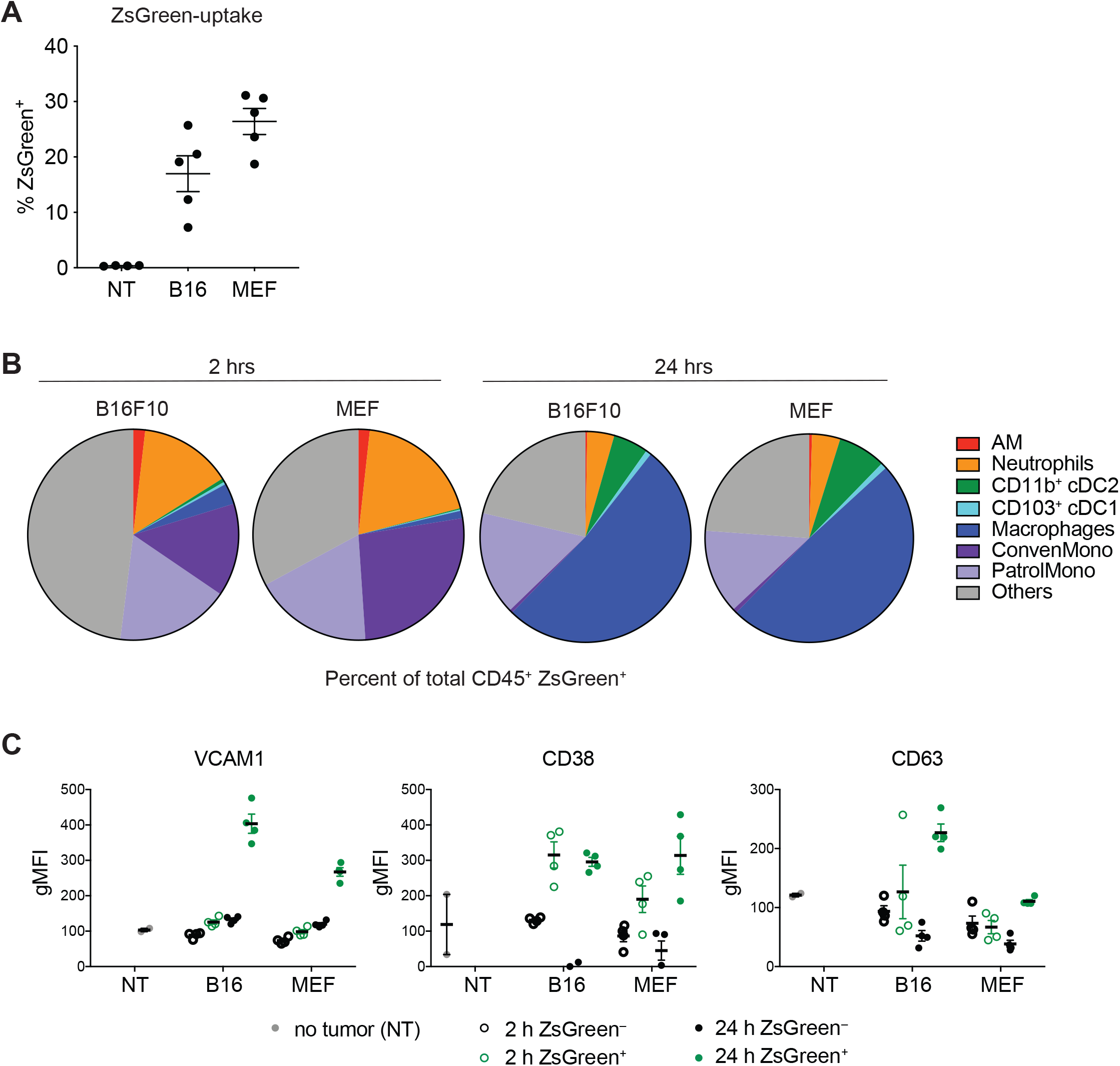
Rapid microparticle-induced macrophage reprogramming is dependent on antigen source. A) Quantification of ZsGreen-uptake by macrophages in the lung 24 hours after i.v. injection with B16ZsGreen (B16), MEFZsGreen (MEF) or non-tumor challenged (NT) mice. B) Quantification of ZsGreen-uptake by distinct myeloid populations in the lung at 2 hours (left) and 24 hours (right) post i.v. injection of B16ZsGreen (B16F10) or MEFZsGreen (MEF) by flow cytometry. C) Quantification of VCAM1, CD38 and CD63 expression on ZsGreen^-^ and ZsGreen^+^ lung macrophages at 2 hours and 24 hours post i.v. injection of B16ZsGreen (B16) or MEFZsGreen (MEF) or non-tumor challenged (NT) mice as determined by flow cytometric analysis.

**Supplemental Figure 4.**
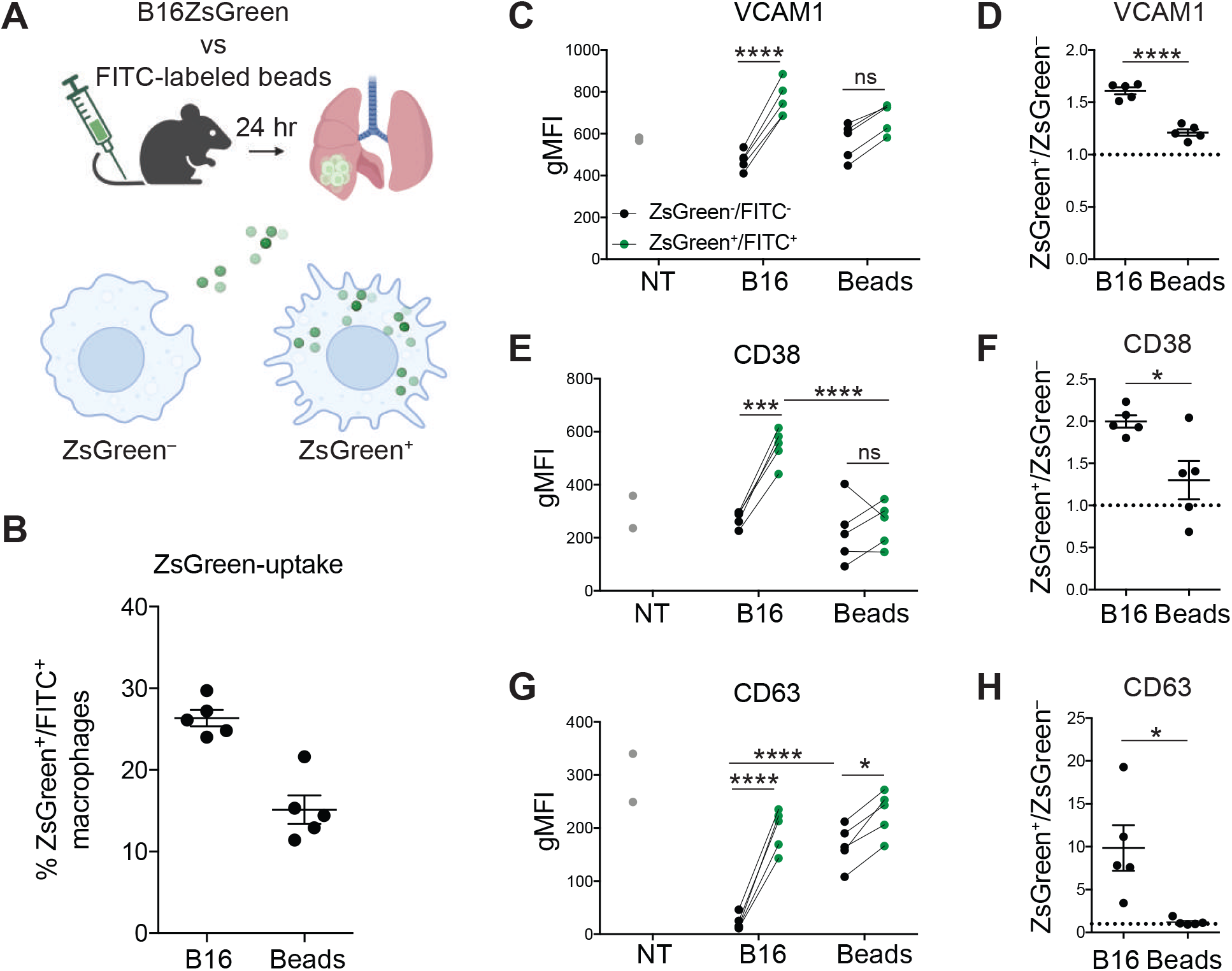
Microparticle-induced macrophage reprogramming is dependent on antigen source. A) Experimental layout. B) Quantification of uptake of microparticles or FITC-labeled beads as ZsGreen^+^ or FITC^+^ lung macrophages 24 hours post i.v. injection of B16ZsGreen or FITC-labeled beads by flow cytometry. C, E, G) Quantification of VCAM1 (C) CD38 (E) and CD63 (G) expression on ZsGreen^-^ and ZsGreen^+^ lung macrophages from indicated groups. D, F, H) Ratio of the expression level of VCAM1 (D), CD38 (F) and CD63 (H) on ZsGreen^+^ over ZsGreen^-^ macrophages. Data are representative of two independent experiments; **** p < 0.0001, *** p < 0.001, ** p < 0.01, * p < 0.05 as determined by the two-way ANOVA and multiple comparison test (C, E, G) or Student’s *t*-test (D, F, H).

**Supplemental Figure 5.**
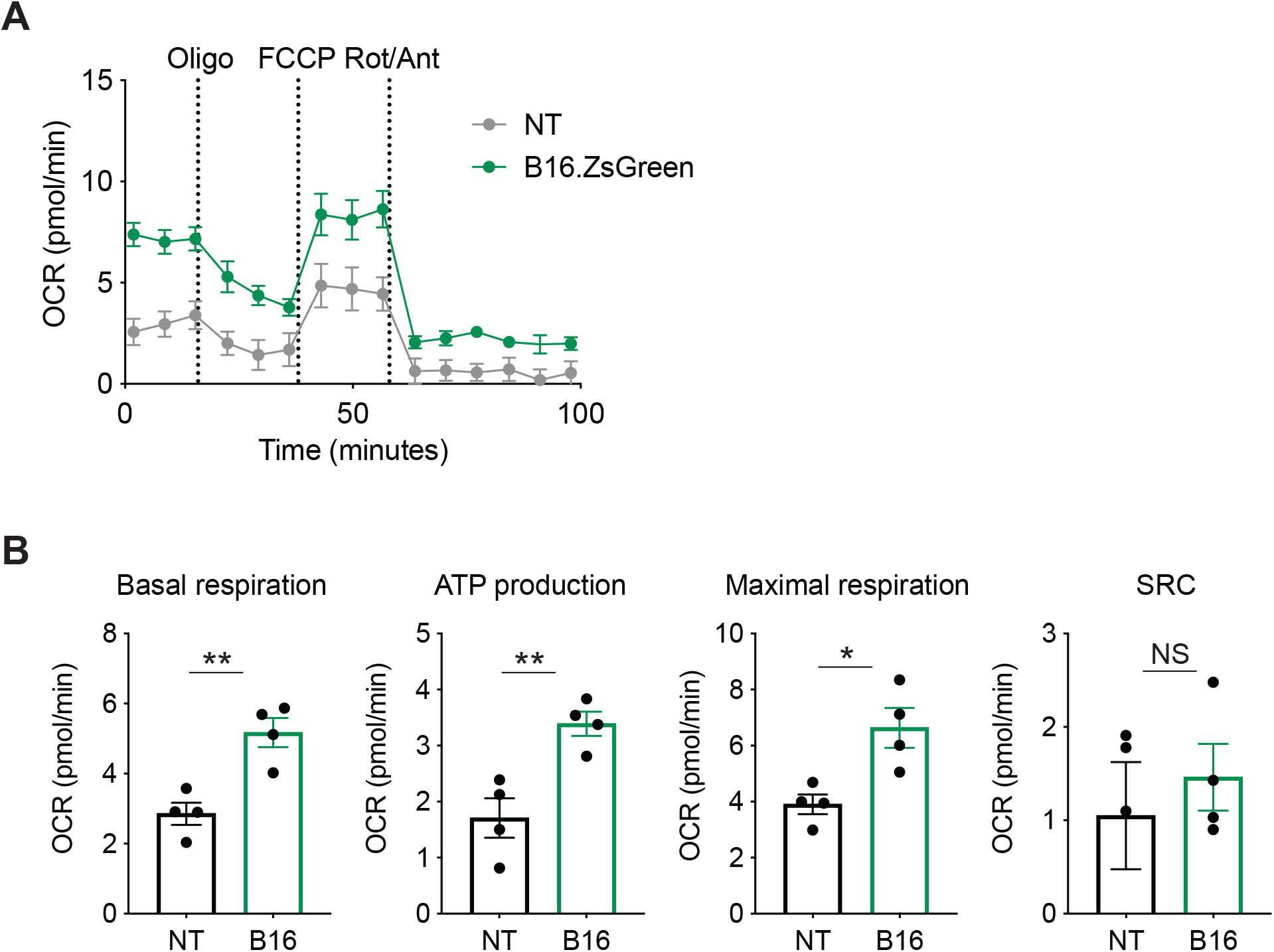
Myeloid cells in the early metastatic lung upregulate mitochondrial oxidative phosphorylation upon encountering B16 melanoma cells. A) CD11b^+^ lung myeloid cells were FACS-sorted from non-tumor challenged (NT) or B16ZsGreen-injected mice and were plated into Seahorse XFe24 plates. Oxygen consumption rate (OCR) in lung myeloid cells was measured by Seahorse assay with Oligomycin (Oligo), FCCP and rotenone and antimycin A (Rot/Ant) added sequentially. B) Quantification of basal respiration, ATP production, maximal respiration, and spare respiratory capacity (SRC) in lung myeloid cells of NT and B16-challenged mice. ** p < 0.01, * p < 0.05 as determined by the Student’s *t*-test.

**Supplemental Figure 6.**
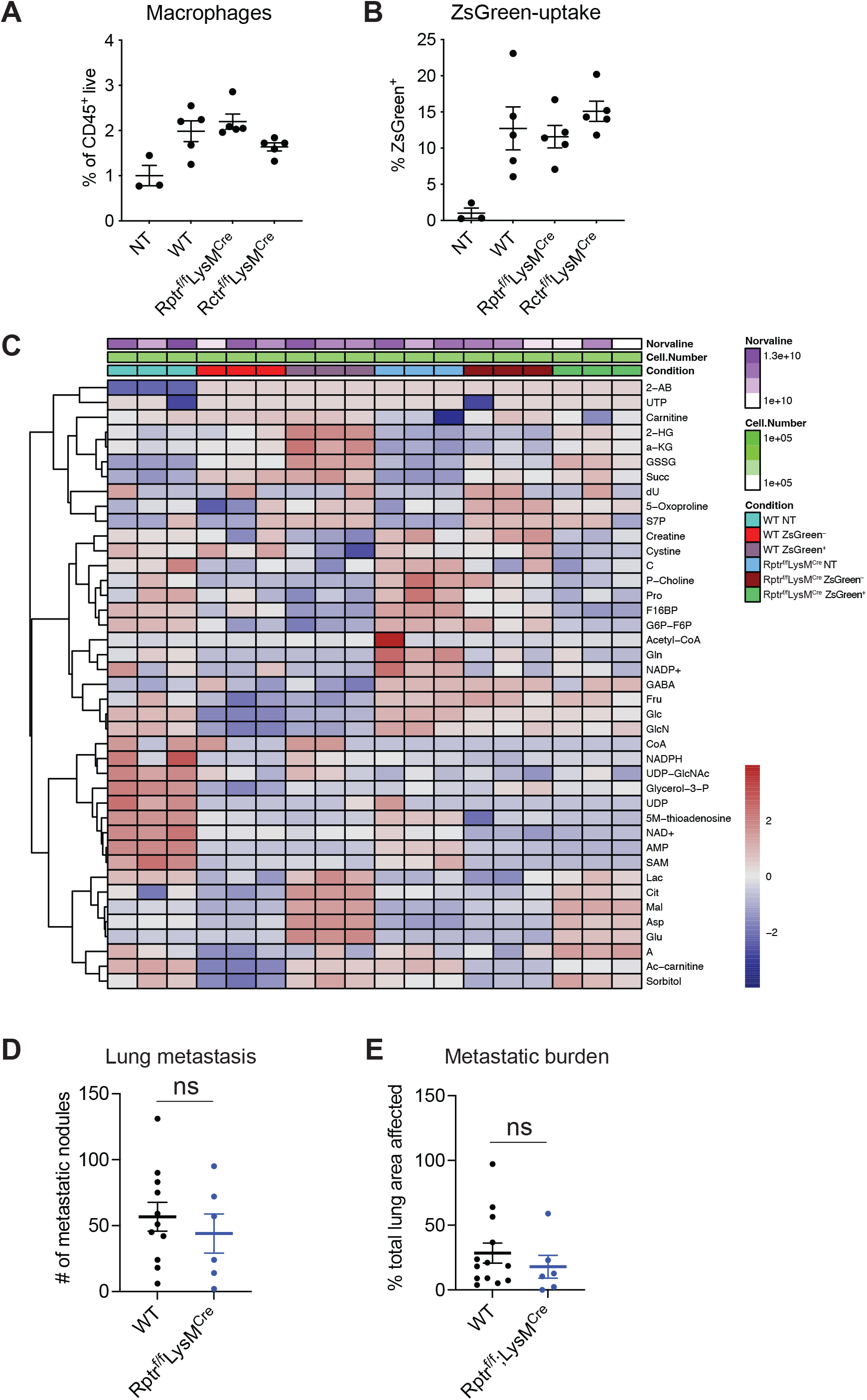
mTORC1-dependent metabolic and phenotypic reprogramming of macrophages in the early metastatic lung. A-B) Quantification of the proportion of macrophages (A) and ZsGreen-uptake (B) by lung macrophages 24 hours post i.v. injection of B16ZsGreen into mTORC1-deficient *Rptr*^*f/f*^*/LysM*^*Cre*^, mTORC2-deficient *Rctr*^*f/f*^*/LysM*^*Cre*^ mice and wild type (WT) littermates as determined by flow cytometry. C) Heatmap of fractional contribution of incorporated ^13^C-glucose in metabolites. WT or *Rptr*^*f/f*^*/LysM*^*Cre*^ BMDM were co-cultured with B16ZsGreen at a 5:1 ratio for 24 hours followed by FACS-sorting of ZsGreen^-^, ZsGreen^+^ and non-tumor challenged (NT) BMDM. Cells were rested for 1 hour and media was substituted with media containing isotopically labeled glucose (^13^C) for 2 hours^19^ upon which cells were fixed and metabolites were extracted for LC/MS analysis. Top two rows represent internal normalization controls (Norvaline and Cell Number). D) Quantification of the number of metastatic nodules in lungs of *Rptr*^*f/f*^*/LysM*^*Cre*^ mice and wild type (WT) littermates on H&E stained microscopic slides of lungs harvested 2-3 weeks after i.v. administration of B16ZsGreen. E) Quantification of the metastatic burden, determined as the area of the total lung lobe affected by metastatic cancer cells based on H&E stained slides. Metastatic nodules and total lung area were manually outlined and ‘area affected’ was calculated using ImageJ.

## STAR Methods

### Resource Availability

### Lead Contact

Further information and requests for resources and reagents should be directed to and will be fulfilled by the Lead Contact, Mark Headley.

## Materials Availability

All unique reagents generated in this study are available from the Lead Contact without restriction.

## Data and Code Availability

Bulk RNA-seq data has been deposited at GEO (xxx) and are publicly available as of the date of publication. Accession numbers are also listed in the Key Resources Table. No original code was written for analyses contained in this manuscript; all analyses were carried out using publicly available resources. DOIs are listed in the Key Resources Table. Any additional information required to reanalyze the data reported in this paper is available from the lead contact upon request.

## Experimental Model and Subject Details

### Mice

All mice were treated in accordance with the regulatory standards of the National Institutes of Health and American Association of Laboratory Animal Care and were approved by the UCSF Institution of Animal Care and Use Committee and were maintained under specific pathogen-free conditions at the University of California, San Francisco Animal Barrier Facility. C57BL/6J mice were purchased from The Jackson Laboratory. *MacBlue* mice were a gift from David Hume (The Roslin Institute). *Rptr*^*f/f*^ and *Rctr*^*f/f*^ mice (a gift from Xin Chen, UCSF) were bred to *LysM*^*Cre*^ mice (strain # 004781, The Jackson Laboratory). MEFs were derived from *Ai6;β-actin*^*Cre*^ mice that were generated by crossing *β-actin*^*Cre*^ (strain # 033984, The Jackson Laboratory) to *Ai6* mice (strain # 007906,The Jackson Laboratory) as previously described^2^. Both male and female mice ranging in age from 6-20 weeks were used for experimentation. Food and water were provided ad libitum.

### Cell lines

B16F10-ZsGreen cells were previously generated in our laboratory as described^2^. B16F1-ZsGreen and Lewis Lung Carcinoma (LLC)-ZsGreen cells were generated through transduction of B16F1 or LLC with empty pSiren-ZsGreen (Clontech). All cells were cultured under standard conditions 37°C in 5% CO2 in DMEM (GIBCO), 10% FCS (Benchmark), 1% Pen/Strep/Glut (Invitrogen) unless stated otherwise.

## Method Details

### Cell line intravenous injections

Adherent tumor cells or MEFs were grown to confluency and harvested using 0.05% Trypsin-EDTA (GIBCO) and washed 3x with PBS (GIBCO). For studies involving acute responses, 5-10×10^5^ cells were injected intravenously (i.v.) via the tail vein in a final volume of 100μl PBS and mice were euthanized 24 hours later. For uptake of beads, 1×10^9^ FITC-labeled polystyrene beads (Sigma) were resuspended in 100μl PBS and injected i.v. and mice were euthanized 24 hours later. For long-term metastasis experiments, 1.5×10^5^ B16ZsGreen cells in PBS were injected i.v. and mice were euthanized 2-3 weeks later and lungs were harvested for analysis.

### Rapamycin treatment in vivo

Rapamycin (Millipore Sigma) at 0.4 mg per kg was delivered intranasally into mice one day prior to and 1 hour post i.v. administration of B16ZsGreen. Mice were euthanized 24 hours later and lungs were harvested for analysis.

### Mouse lung digestion

Lungs were collected from mice following euthanasia by overdose with 2.5% Avertin. Lungs were placed in 3 ml of DMEM (GIBCO) in C-Tubes (Miltenyi) briefly processed with a GentleMACS Dissociator (Miltenyi). 2 ml of DMEM with 0.26 U ml^−1^ LiberaseTM (Roche) and 0.25 mg ml^−1^ DNase I (Roche) was subsequently added and samples were then incubated at 37°C in a shaker for 30 min and dissociated to single cell suspensions by GentleMACS. Tissue homogenate was then passed through a 100 μm filter. Red blood cells were lysed with 3 ml of 175 mM NH4Cl per lung for 5 min at room temperature. Samples were then washed with FACS buffer (2% FBS in PBS) and resuspended in appropriate buffer for staining for flow cytometry or FACS-sorting.

### Flow cytometry and Fluorescence Activated Cell Sorting

For flow cytometric analyses, cells were washed with PBS prior to staining with Zombie NIR Fixable live/dead dye (Biolegend) for 20 min at 4°C. Cells were washed in PBS followed by surface staining for 30 min at 4°C with directly conjugated antibodies diluted in FACS buffer containing anti-CD16/32 (clone 2.4G2; BioXCell) to block non-specific binding. Cells were washed again with FACS buffer. For intracellular staining, cells were fixed for 20 min at 4°C using the FOXP3 Fix/Perm kit (BD Biosciences), and washed in permeabilization buffer. Antibodies against intracellular targets were diluted in permeabilization buffer and cells were incubated for 30 min at 4°C followed by another wash prior to read-out on a BD LSR Fortessa SORP cytometer.

For FACS-sorting, cells were washed in PBS followed by surface staining for 30 min at 4°C with directly conjugated antibodies diluted in FACS buffer containing anti-CD16/32. Cells were washed with FACS buffer and filtered over a 70μm mesh. Immediately prior to sorting, DAPI was added to exclude dead cells. Cells were sorted on a BD FACSAria Fusion and BD FACSAria2. Sorted myeloid cells were collected in DMEM (GIBCO), 10% FCS (Benchmark), Pen/Strep/Glut (Invitrogen) at 4°C for further use ex vivo.

For mitochondrial analysis, Mitotracker Deep Red FM (Invitrogen) at 200nM and TMRM (Thermo Fisher Scientific) at 100nM were used for staining of lung cells for the detection of mitochondrial mass and membrane potential. Cells were stained with Mitotracker and TMRM in DMEM at 37°C for 30 mins before staining with surface antibodies.

### RNA sequencing

ZsGreen^+^ and ZsGreen^-^ macrophages were isolated from lungs of mice 24 hours post i.v. injection with B16ZsGreen. Cells were double sorted to ensure high purity. 200 cells were isolated for each sample with 4 biological replicates. Cells were sorted directly into 100ul of Arcturus PicoPure Lysis Buffer. RNA was prepared using the Arcturus PicoPure RNA Isolation Kit (ThermoFisher). Libraries were prepared for each sample utilizing the Nugen Ovation Library Preparation Kit (Tecan). Single End 50bp sequencing was performed on an Illumina HiSeq 3000. Data were aligned to the *Mus musculus* Ensembl GRCm38 v.78 genome and those reads uniquely mapping to Ensembl IDs were tabulated using STAR_2.4.2a. The resulting sample by gene matrix was passed to DESeq2 for downstream normalization and differential expression testing.

### Pathway Enrichment Analysis

For gene set enrichment analysis of samples derived from ZsGreen^+^ and ZsGreen^-^ macrophages, log2 fold changes between groups were calculated for each protein coding gene that was expressed at greater than 5 counts per million (cpm). The resulting, ordered list was passed to the GSEApreRanked function, implemented in GSEA software, downloaded from the Broad Institute (www.gsea-msigdb.org/gsea/index.jsp). GSEApreRanked was run with default parameters on both Hallmark and c5 gene set collections contained in the Broad’s Molecular Signatures Database (MSigDB). Results from the c5 gene set were passed to Cytoscape (www.cytoscape.org) software (v3.71) which was run using default parameters to generate pathway network plots.

### Infinity Flow

Infinity Flow was performed as previously described^9^. Briefly, single cell suspensions of mouse lungs were washed with PBS prior to staining with Zombie NIR Fixable live/dead dye (Biolegend) for 20 min at room temperature (RT). Following staining, a 10-fold volume of Cell Staining Buffer (BioLegend) was added to neutralize any unbound dye and cells were centrifuged at 300*g* for 5 min. Cells were resuspended at 20 × 10^6^ cells/ml in Cell Staining Buffer, and nonspecific staining was then blocked by addition of anti-CD16/32 (2 μg/ml; mouse TrueStain FcX, BioLegend), 2% rat serum (Invitrogen), and 2% Armenian hamster serum (Innovative Research) followed by 15 min incubation at 4°C. Cells were then washed, resuspended in Cell Staining Buffer and a master mix of the indicated Backbone antibody panel^9^ at 20 × 10^6^ cells/ml, and incubated for 30 min at 4°C. Following Backbone staining, cells were washed twice and resuspended at 20 × 10^6^ cells/ml in Cell Staining Buffer and 75 μl was added to each well of the LEGENDScreen plates. Staining and fixation were performed exactly as per manufacturer directions. Not a portion of the dataset described, here in Figure 1, was published in ^9^ alongside the Infinity Flow method. This data represents an expanded dataset and the analysis is non-overlapping.

### Two photon imaging of mouse lung slices and analysis

Slice imaging was performed as previously described^2^. Briefly, *MacBlue* mice were injected with 1×10^6^ B16ZsGreen cells through the tail vein. After 24 hours, mice were euthanized by anesthetic overdose with 1 ml 2.5% Avertin and then intubated by tracheotomy with the sheath from an 18-gauge i.v. catheter. Lungs were subsequently inflated with 1 ml of 2% low melting agarose (BMA) in sterile PBS at 37°C. Agarose was then solidified by flooding the chest cavity with 4°C PBS. Inflated lungs were excised, and the left lobe was cut into 250μm sections using a vibratome. Sections were mounted on plastic coverslips and imaged by two-photon microscopy at 37°C in carbogen (5% CO2:95% O2)-perfused RPMI-1640 media (GIBCO, without Phenol Red) in a heated chamber. The Maitai laser of two-photon microscope was set to 800nm for excitation of CFP. The Chameleon laser was set to 950nm for excitation of ZsGreen. Emitted light was detected using a 25x 1.2NA water lens (Zeiss) coupled to a 6-color detector array (custom; using Hamamatsu H9433MOD detectors). Emission filters used were blue 475/23, green 510/42, yellow 542/27, red 607/70, far red 675/67. The microscope was controlled by the MicroManager software suite, and time-lapse z-stack images were acquired every 90 seconds with five-fold averaging and z-step of 4μm. Data analysis was performed with Imaris software (Bitplane).

### Bone marrow-derived macrophage (BMDM) and B16ZsGreen coculture

Bone marrow was obtained from femurs and tibia of C57BL6/J mice and cultured in DMEM (GIBCO), 10% FCS (Benchmark), Pen/Strep/Glut (Invitrogen) in the presence of 20ng/ml M-CSF (Peprotech) for 5 days. BMDM were then co-cultured with B16ZsGreen cells at a 5:1 ratio for 24 hours before assays.

### Extracellular Flux Analysis

Extracellular flux was measured using a Seahorse XFe24 or XFe96 Analyzer (Agilent) using Mito Stress Test Kits (Aligent Technologies) per manufacturer instructions. Sorted CD11b^+^ BMDM or lung macrophages were plated in poly-L-lysine-coated XFe24 or XFe96 plates for one hour before the assay. Respiration was measured in the basal state and in response to 1μM oligomycin, 1μM FCCP and 0.5μM rotenone and antimycin A.

### Quantification of ATP luminescence

ATP production from FACS-sorted lung macrophages was measured using Luminescent ATP Detection Assay Kit (Abcam) per manufacturer instructions. Lung macrophages were plated in poly-L-lysine-coated 96 flat bottom plates one hour prior to ATP detection. Luminescence was measured using FlexStation 3 Multi-Mode Microplate Reader (Molecular Devices).

### Metabolic tracing of isotopically labeled ^13^C-glucose

BMDM and B16ZsGreen co-culture was performed as noted above. After 24 hours, cells were collected and FACS-sorted based on ZsGreen-positivity (ZsGreen^−^, ZsGreen^+^ and non-tumor challenged (NT)). X cells were cultured in xx flat bottom plates and allowed to rest for 1 hour in D10. The medium was exchanged for DMEM medium containing 25 mM ^13^C-glucose (Cambridge Isotope Laboratories Inc., CLM-1396) and no additional ^12^C-glucose for 2 hours. Cells were washed twice with PBS and extracted with mass spectrometry grade 80% methanol (ThermoFisher, A456-1) and 20% water (ThermoFisher, W6500) supplemented with 5 nmol DL-Norvaline (Sigma, N7502). Protein concentrations of the methanol extract were determined via BCA (Pierce, 23225) with no significant variability assessed (5 μL transferred into 45 μL RIPA buffer, 5 μL of the RIPA dissolved solution assayed). Insoluble material was pelleted in a 4ºC centrifuge at 16,000g, the supernatant was transferred and dried in a Speedvac. Dried metabolites were resuspended in 50% ACN:water and 1/10th of the volume was loaded onto a Luna 3 um NH2 100A (150 × 2.0 mm) column (Phenomenex). The chromatographic separation was performed on a Vanquish Flex (Thermo Scientific) with mobile phases A (5 mM NH_4_AcO pH 9.9) and B (ACN) and a flow rate of 200 μL/min. A linear gradient from 15% A to 95% A over 18 min was followed by 9 min isocratic flow at 95% A and re-equilibration to 15% A. Metabolites were detected with a Thermo Scientific Q Exactive mass spectrometer run with polarity switching (+3.5 kV / -3.5 kV) in full scan mode with an m/z range of 65-975. TraceFinder 4.1 (Thermo Scientific) was used to quantify the targeted metabolites by area under the curve using expected retention time and accurate mass measurements (< 5 ppm). Values were normalized to cell number and sample protein concentration. Relative amounts of metabolites were calculated by summing up the values for all isotopologues of a given metabolite. Fractional contribution (FC) of ^13^C carbons to total carbon for each metabolite was calculated ^34^. Data analysis was performed using in-house developed R scripts.

### Quantification of metastatic burden

Metastatic seeding: The proportion of CD45^−^ZsGreen^+^ cells was gated of total live cells from total lung single cell suspensions 24 hours after i.v. administration of 5-10×10^5^ B16ZsGreen cells using flow cytometry.

Overt metastatic disease: The number of metastatic foci on the surface of all the lobes were counted by eye by two blinded researchers under a dissection microscope. The left lobe was fixed in 4% PFA and processed into formalin-fixed paraffin-embedded tissue blocks. One section was stained in H&E and the total area of the lung affected by metastasis was quantified by manually outlining metastatic nodules and total lung lobe in ImageJ. The metastatic burden was calculated as the percentage of area affected by metastatic nodules as a fraction of the total lung lobe area.

## Graphical Abstract

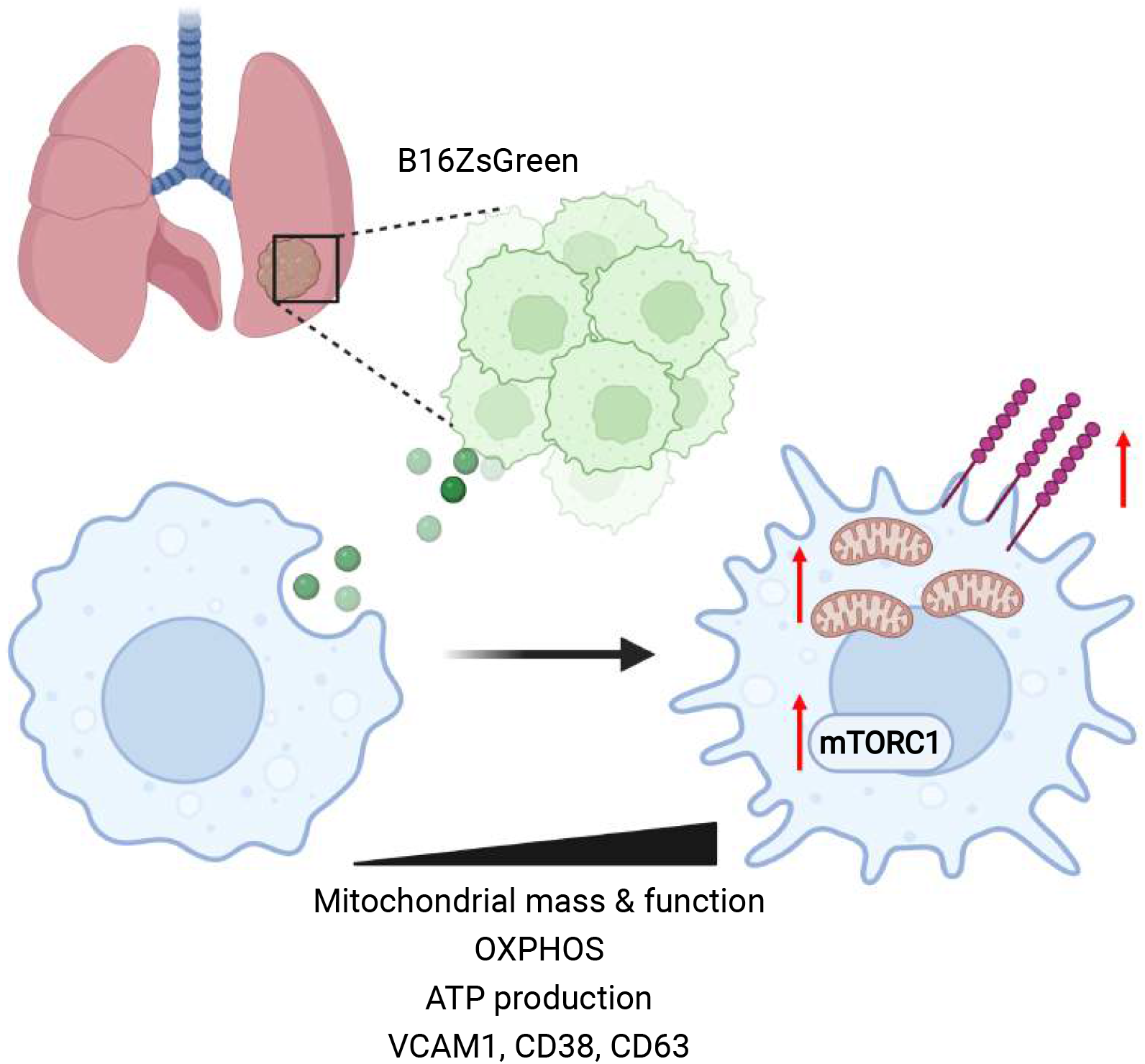

## References

1. Kitamura, T., Qian, B.-Z., and Pollard, J.W. (2015). Immune cell promotion of metastasis. Nat. Rev. Immunol. 15, 73–86. 10.1038/nri3789.

2. Headley, M.B., Bins, A., Nip, A., Roberts, E.W., Looney, M.R., Gerard, A., and Krummel, M.F. (2016). Visualization of immediate immune responses to pioneer metastatic cells in the lung. Nature 531, 513–517. 10.1038/nature16985.

3. DeNardo, D.G., and Ruffell, B. (2019). Macrophages as regulators of tumour immunity and immunotherapy. Nat. Rev. Immunol. 19, 369–382. 10.1038/s41577-019-0127-6.

4. Hanna, R.N., Cekic, C., Sag, D., Tacke, R., Thomas, G.D., Nowyhed, H., Herrley, E., Rasquinha, N., McArdle, S., Wu, R., et al. (2015). Patrolling monocytes control tumor metastasis to the lung. Science 350, 985–990. 10.1126/science.aac9407.

5. Morrissey, S.M., Zhang, F., Ding, C., Montoya-Durango, D.E., Hu, X., Yang, C., Wang, Z., Yuan, F., Fox, M., Zhang, H.-G., et al. (2021). Tumor-derived exosomes drive immunosuppressive macrophages in a pre-metastatic niche through glycolytic dominant metabolic reprogramming. Cell Metab. 33, 2040-2058.e10. 10.1016/j.cmet.2021.09.002.

6. Costa-Silva, B., Aiello, N.M., Ocean, A.J., Singh, S., Zhang, H., Thakur, B.K., Becker, A., Hoshino, A., Mark, M.T., Molina, H., et al. (2015). Pancreatic cancer exosomes initiate pre-metastatic niche formation in the liver. Nat. Cell Biol. 17, 816–826. 10.1038/ncb3169.

7. Hoshino, A., Costa-Silva, B., Shen, T.-L., Rodrigues, G., Hashimoto, A., Tesic Mark, M., Molina, H., Kohsaka, S., Di Giannatale, A., Ceder, S., et al. (2015). Tumour exosome integrins determine organotropic metastasis. Nature 527, 329–335. 10.1038/nature15756.

8. Peinado, H., Alečkovic, M., Lavotshkin, S., Matei, I., Costa-Silva, B., Moreno-Bueno, G., Hergueta-Redondo, M., Williams, C., García-Santos, G., Ghajar, C., et al. (2012). Melanoma exosomes educate bone marrow progenitor cells toward a pro-metastatic phenotype through MET. Nat. Med. 18, 883–891. 10.1038/nm.2753.

9. Becht, E., Tolstrup, D., Dutertre, C.-A., Morawski, P.A., Campbell, D.J., Ginhoux, F., Newell, E.W., Gottardo, R., and Headley, M.B. (2021). High-throughput single-cell quantification of hundreds of proteins using conventional flow cytometry and machine learning. Sci. Adv. 7, eabg0505. 10.1126/sciadv.abg0505.

10. Barry, K.C., Hsu, J., Broz, M.L., Cueto, F.J., Binnewies, M., Combes, A.J., Nelson, A.E., Loo, K., Kumar, R., Rosenblum, M.D., et al. (2018). A natural killer-dendritic cell axis defines checkpoint therapy-responsive tumor microenvironments. Nat. Med. 24, 1178–1191. 10.1038/s41591-018-0085-8.

11. Broz, M.L., Binnewies, M., Boldajipour, B., Nelson, A.E., Pollack, J.L., Erle, D.J., Barczak, A., Rosenblum, M.D., Daud, A., Barber, D.L., et al. (2014). Dissecting the Tumor Myeloid Compartment Reveals Rare Activating Antigen-Presenting Cells Critical for T Cell Immunity. Cancer Cell 26, 638–652. 10.1016/j.ccell.2014.09.007.

12. Ovchinnikov, D.A., van Zuylen, W.J.M., DeBats, C.E.E., Alexander, K.A., Kellie, S., and Hume, D.A. (2008). Expression of Gal4-dependent transgenes in cells of the mononuclear phagocyte system labeled with enhanced cyan fluorescent protein using Csf1r-Gal4VP16/UAS-ECFP double-transgenic mice. J. Leukoc. Biol. 83, 430–433. 10.1189/jlb.0807585.

13. Cunningham, J.T., Rodgers, J.T., Arlow, D.H., Vazquez, F., Mootha, V.K., and Puigserver, P. (2007). mTOR controls mitochondrial oxidative function through a YY1-PGC-1alpha transcriptional complex. Nature 450, 736–740. 10.1038/nature06322.

14. Morita, M., Gravel, S.-P., Chénard, V., Sikström, K., Zheng, L., Alain, T., Gandin, V., Avizonis, D., Arguello, M., Zakaria, C., et al. (2013). mTORC1 controls mitochondrial activity and biogenesis through 4E-BP-dependent translational regulation. Cell Metab. 18, 698–711. 10.1016/j.cmet.2013.10.001.

15. Covarrubias, A.J., Aksoylar, H.I., Yu, J., Snyder, N.W., Worth, A.J., Iyer, S.S., Wang, J., Ben-Sahra, I., Byles, V., Polynne-Stapornkul, T., et al. (2016). Akt-mTORC1 signaling regulates Acly to integrate metabolic input to control of macrophage activation. eLife 5. 10.7554/eLife.11612.

16. Huang, S.C.-C., Smith, A.M., Everts, B., Colonna, M., Pearce, E.L., Schilling, J.D., and Pearce, E.J. (2016). Metabolic Reprogramming Mediated by the mTORC2-IRF4 Signaling Axis Is Essential for Macrophage Alternative Activation. Immunity 45, 817–830. 10.1016/j.immuni.2016.09.016.

17. Hallowell, R.W., Collins, S.L., Craig, J.M., Zhang, Y., Oh, M., Illei, P.B., Chan-Li, Y., Vigeland, C.L., Mitzner, W., Scott, A.L., et al. (2017). mTORC2 signalling regulates M2 macrophage differentiation in response to helminth infection and adaptive thermogenesis. Nat. Commun. 8, 14208. 10.1038/ncomms14208.

18. Weichhart, T., Hengstschläger, M., and Linke, M. (2015). Regulation of innate immune cell function by mTOR. Nat. Rev. Immunol. 15, 599–614. 10.1038/nri3901.

19. Jang, C., Chen, L., and Rabinowitz, J.D. (2018). Metabolomics and Isotope Tracing. Cell 173, 822–837. 10.1016/j.cell.2018.03.055.

20. Joyce, J.A., and Pollard, J.W. (2009). Microenvironmental regulation of metastasis. Nat. Rev. Cancer 9, 239–252. 10.1038/nrc2618.

21. Güç, E., and Pollard, J.W. (2021). Redefining macrophage and neutrophil biology in the metastatic cascade. Immunity 54, 885–902. 10.1016/j.immuni.2021.03.022.

22. Gener Lahav, T., Adler, O., Zait, Y., Shani, O., Amer, M., Doron, H., Abramovitz, L., Yofe, I., Cohen, N., and Erez, N. (2019). Melanoma-derived extracellular vesicles instigate proinflammatory signaling in the metastatic microenvironment. Int. J. Cancer 145, 2521–2534. 10.1002/ijc.32521.

23. Van den Bossche, J., O’Neill, L.A., and Menon, D. (2017). Macrophage Immunometabolism: Where Are We (Going)? Trends Immunol. 38, 395–406. 10.1016/j.it.2017.03.001.

24. Mazzone, M., Menga, A., and Castegna, A. (2018). Metabolism and TAM functions-it takes two to tango. FEBS J. 285, 700–716. 10.1111/febs.14295.

25. Vitale, I., Manic, G., Coussens, L.M., Kroemer, G., and Galluzzi, L. (2019). Macrophages and Metabolism in the Tumor Microenvironment. Cell Metab. 30, 36–50. 10.1016/j.cmet.2019.06.001.

26. Argüello, R.J., Combes, A.J., Char, R., Gigan, J.-P., Baaziz, A.I., Bousiquot, E., Camosseto, V., Samad, B., Tsui, J., Yan, P., et al. (2020). SCENITH: A Flow Cytometry-Based Method to Functionally Profile Energy Metabolism with Single-Cell Resolution. Cell Metab. 32, 1063–1075. 10.1016/j.cmet.2020.11.007.

27. Li, S., Yu, J., Huber, A., Kryczek, I., Wang, Z., Jiang, L., Li, X., Du, W., Li, G., Wei, S., et al. (2022). Metabolism drives macrophage heterogeneity in the tumor microenvironment. Cell Rep. 39, 110609. 10.1016/j.celrep.2022.110609.

28. Artyomov, M.N., Sergushichev, A., and Schilling, J.D. (2016). Integrating immunometabolism and macrophage diversity. Semin. Immunol. 28, 417–424. 10.1016/j.smim.2016.10.004.

29. Laviron, M., Petit, M., Weber-Delacroix, E., Combes, A.J., Arkal, A.R., Barthélémy, S., Courau, T., Hume, D.A., Combadière, C., Krummel, M.F., et al. (2022). Tumor-associated macrophage heterogeneity is driven by tissue territories in breast cancer. Cell Rep. 39, 110865. 10.1016/j.celrep.2022.110865.

30. Mujal, A.M., Combes, A.J., Rao, A.A., Binnewies, M., Samad, B., Tsui, J., Boissonnas, A., Pollack, J.L., Argüello, R.J., Meng, M.V., et al. (2022). Holistic Characterization of Tumor Monocyte-to-Macrophage Differentiation Integrates Distinct Immune Phenotypes in Kidney Cancer. Cancer Immunol. Res., canimm.0588.2021. 10.1158/2326-6066.CIR-21-0588.

31. Chen, W., Ma, T., Shen, X., Xia, X., Xu, G., Bai, X., and Liang, T. (2012). Macrophage-induced tumor angiogenesis is regulated by the TSC2-mTOR pathway. Cancer Res. 72, 1363–1372. 10.1158/0008-5472.CAN-11-2684.

32. Wenes, M., Shang, M., Di Matteo, M., Goveia, J., Martín-Pérez, R., Serneels, J., Prenen, H., Ghesquière, B., Carmeliet, P., and Mazzone, M. (2016). Macrophage Metabolism Controls Tumor Blood Vessel Morphogenesis and Metastasis. Cell Metab. 24, 701–715. 10.1016/j.cmet.2016.09.008.

33. Pittet, M.J., Michielin, O., and Migliorini, D. (2022). Clinical relevance of tumour-associated macrophages. Nat. Rev. Clin. Oncol. 19, 402–421. 10.1038/s41571-022-00620-6.

34. Cameron, A.M., Castoldi, A., Sanin, D.E., Flachsmann, L.J., Field, C.S., Puleston, D.J., Kyle, R.L., Patterson, A.E., Hässler, F., Buescher, J.M., et al. (2019). Inflammatory macrophage dependence on NAD+ salvage is a consequence of reactive oxygen species-mediated DNA damage. Nat. Immunol. 20, 420–432. 10.1038/s41590-019-0336-y.

