## Supplementary material for "Uptake of tumor-derived microparticles induces metabolic reprogramming of macrophages in the early metastatic lung": Key Resources

| Reagent or Resource | Source | Identifier |
|---|---|---|
| *Antibodies* | | |
| anti-mouse CD11b - BV605 (clone M1/70) | Biolegend | 101237 |
| anti-mouse CD11c - BV510 (clone N418) | Biolegend | 117337 |
| anti-mouse CD11c - PerCp-Cy5.5 (clone N418) | Biolegend | 117328 |
| anti-mouse CD19 - BV785 (clone 6D5) | Biolegend | 115543 |
| anti-mouse CD24 - PECy7 (clone M1/69) | Biolegend | 101822 |
| anti-mouse CD38 - BV711 (clone 90/CD38) | BD BioSciences | 740697 |
| anti-mouse CD38 - PE (clone 14.27) | Biolegend | 250505 |
| anti-mouse CD45 - PerCp-Cy5.5 (clone 30-F11) | Biolegend | 103132 |
| anti-mouse CD45 - AF700 (clone 30-F11) | Biolegend | 103128 |
| anti-mouse CD45R (B220) - BV785 (clone RA3-6B2) | Biolegend | 103245 |
| anti-mouse CD63 - PE (clone NVG-2) | Biolegend | 143904 |
| anti-mouse CD90.2 - BV785 (clone 30-H12) | Biolegend | 105331 |
| anti-mouse CD103 - APC (clone 2E7) | Biolegend | 121414 |
| anti-mouse CD106 - BV650 (clone 429(MVCAM.A)) | BD BioSciences | 740471 |
| anti-mouse CD106 - PE (clone 429 (MVCAM.A)) | Biolegend | 105713 |
| anti-mouse Ly6C - BV711 (clone HK1.4) | Biolegend | 128037 |
| anti-mouse Ly6G - BV785 (clone 1A8) | Biolegend | 127645 |
| anti-mouse MHCII (I-A/I-E) - AF700 (clone M5/114.15.2) | Biolegend | 107622 |
| anti-mouse MHCII (I-A/I-E) - BV650 (clone M5/114.15.2) | Biolegend | 107641 |
| anti-mouse NK1.1 - BV785 (clone PK136) | Biolegend | 108749 |
| anti-mouse SiglecF - BV785 (clone E50-2440) | BD BioSciences | 740956 |
| Phospho-S6 Ribosomal Protein (ser235/236)XP - PE (clone [ | Cell Signaling Technologies | 5316S |
| Phospho-4E-BP1(Thr37/46)- AF647 (clone 236B4) | Cell Signaling Technologies | 5123S |
| anti-mouse CD16/32 (clone 2.4G2) | BioXCell | BE0307 |
| Normal Rat Serum | Thermo Fisher | 10710C |
| *Biological Samples* | | |
| Mouse tssue samples (LN, tumor) | UC San Francisco | IACUC: AN184232 |
| *Chemicals, Peptides, and Recombinant Proteins* | | |
| Dnase I | Millipore Sigma | 10104159001 |
| Liberase TM | Roche | 5401127001 |
| Rapamycin | Millipore Sigma | 553210 |
| Zombie NIR Fixable Viability Dye | Biolegend | 423106 |
| Mitotracker Deep Red FM | Thermo Fisher Scientific | M22426 |
| TMRM | Thermo Fisher Scientific | I34361 |
| Low Melting Agarose | BMA | |
| Recombinant murine M-CSF | Peprotech | 315-02 |
| poly-L-lysine | | |
| *Critical Commercial Assays* | | |
| Foxp3/ Transcription Factor Staining Buffer Kit | BD Biosciences | 554655 |
| Seahorse XFe24 Analyzer Extracellular Flux Assay Kit | Agilent Technologies | 102340-100 |
| Seahorse XFe96 Analyzer Extracellular Flux Assay Kit | Agilent Technologies | 102416-100 |
| Seahorse XF Cell Mito Stress Test Kit | Agilent Technologies | 103015-100 |
| UltraComp eBeads Compensation Beads | Fisher Scientific | 01-2222-42 |
| Luminescent ATP Detection Assay Kit | Abcam | ab113849 |
| *Experimental Models: Cell lines* | | |
| B16F1 | | |
| B16F10 | AATCC | CRL-6475 |
| B16ZsGreen | UC San Francisco | N/A |
| LLC | | |
| MEF | | |
| *Experimental Models: Organisms/Strains* | | |
| Mouse: C57BL/6J | The Jackson Laboratory | Stock # 000664 |
| Mouse: MacBlue | David Hume, Roslin Institute | |
| Mouse: Raptor F/F | Chen, University of California San Francisco | |
| Mouse: Rictor F/F | Chen, University of California San Francisco | |
| Mouse: LysM-Cre (B6.129P2-Lyz2tm1(cre)Ifo/J) | The Jackson Laboratory | Stock # 004781 |
| Mouse: β-actin-Cre (B6.FVB-Tmem163Tg(ACTB-cre)2Mrt/Em | The Jackson Laboratory | Stock # 033984 |
| Mouse: Ai6 (B6.Cg-Gt(ROSA)26Sortm6(CAG-ZsGreen1)Hze/ | The Jackson Laboratory | Stock # 007906 |
| *Deposited Data* | | |
| *Software and Algorithms* | | |
| Imaris | Bitplane | https://imaris.exinst.com/ |
| ImageJ | NIH | https://imagej.nih.gov/ij/ |
| FlowJo | Becton Dickinson | https://flowjo.com/ |
| MicroManager | | |
