## Supplementary material for "Uptake of tumor-derived microparticles induces metabolic reprogramming of macrophages in the early metastatic lung": Table S1

| Cluster | Pathway |
|---|---|
| | **Translation** |
| A | GO_COFACTOR_METABOLIC_PROCESS |
| A | GO_RIBONUCLEOPROTEIN_COMPLEX |
| A | GO_NCRNA_PROCESSING |
| A | GO_COFACTOR_BIOSYNTHETIC_PROCESS |
| A | GO_NCRNA_METABOLIC_PROCESS |
| A | GO_TRNA_METABOLIC_PROCESS |
| A | GO_RNA_PHOSPHODIESTER_BOND_HYDROLYSIS |
| A | GO_MATURATION_OF_5_8S_RRNA |
| A | GO_RIBOSOME_BIOGENESIS |
| A | GO_RRNA_METABOLIC_PROCESS |
| A | GO_RNA_PROCESSING |
| A | GO_MRNA_METABOLIC_PROCESS |
| A | GO_RIBONUCLEOPROTEIN_COMPLEX_BIOGENESIS |
| A | GO_QUINONE_METABOLIC_PROCESS |
| A | GO_COENZYME_METABOLIC_PROCESS |
| | **Protein processing** |
| B | GO_ANTIGEN_PROCESSING_AND_PRESENTATION_OF_PEPTIDE_ANTIGEN_VIA_MHC_CLASS_I |
| B | GO_PROTEASOMAL_PROTEIN_CATABOLIC_PROCESS |
| B | GO_MACROMOLECULE_CATABOLIC_PROCESS |
| B | GO_PROTEIN_MODIFICATION_BY_SMALL_PROTEIN_CONJUGATION_OR_REMOVAL |
| B | GO_ERAD_PATHWAY |
| B | GO_PROTEIN_CATABOLIC_PROCESS |
| B | GO_ER_ASSOCIATED_UBIQUITIN_DEPENDENT_PROTEIN_CATABOLIC_PROCESS |
| | **Catabolic processes** |
| C | GO_LIPID_CATABOLIC_PROCESS |
| C | GO_MICROBODY_PART |
| C | GO_FATTY_ACID_BETA_OXIDATION |
| C | GO_MICROBODY |
| C | GO_ORGANIC_ACID_CATABOLIC_PROCESS |
| C | GO_CELLULAR_LIPID_CATABOLIC_PROCESS |
| C | GO_GLYCOSYL_COMPOUND_CATABOLIC_PROCESS |
| C | GO_FATTY_ACID_CATABOLIC_PROCESS |
| C | GO_LIPID_METABOLIC_PROCESS |
| C | GO_SINGLE_ORGANISM_CATABOLIC_PROCESS |
| C | GO_SMALL_MOLECULE_CATABOLIC_PROCESS |
| C | GO_ORGANIC_ACID_METABOLIC_PROCESS |
| C | GO_MICROBODY_LUMEN |
| C | GO_FATTY_ACID_METABOLIC_PROCESS |
| C | GO_MONOCARBOXYLIC_ACID_METABOLIC_PROCESS |
| C | GO_SULFUR_COMPOUND_BIOSYNTHETIC_PROCESS |
| | **Mitochondrial respiration** |
| D | GO_MITOCHONDRIAL_TRANSLATION |
| D | GO_NUCLEOBASE_CONTAINING_SMALL_MOLECULE_METABOLIC_PROCESS |
| D | GO_ORGANOPHOSPHATE_METABOLIC_PROCESS |
| D | GO_INTRINSIC_COMPONENT_OF_MITOCHONDRIAL_MEMBRANE |
| D | GO_GPI_ANCHOR_METABOLIC_PROCESS |
| D | GO_ORGANELLE_INNER_MEMBRANE |
| D | GO_ENERGY_DERIVATION_BY_OXIDATION_OF_ORGANIC_COMPOUNDS |
| D | GO_LIPOPROTEIN_METABOLIC_PROCESS |
| D | GO_ORGANELLAR_RIBOSOME |
| D | GO_CELLULAR_RESPIRATION |
| D | GO_GENERATION_OF_PRECURSOR_METABOLITES_AND_ENERGY |
| D | GO_OXIDATIVE_PHOSPHORYLATION |
| D | GO_OXIDOREDUCTASE_ACTIVITY_ACTING_ON_NAD_P_H_QUINONE_OR_SIMILAR_COMPOUND_AS_ACCEPTOR |
| D | GO_MITOCHONDRIAL_ENVELOPE |
| D | GO_MITOCHONDRIAL_RESPIRATORY_CHAIN_COMPLEX_ASSEMBLY |
| D | GO_CELLULAR_PROTEIN_COMPLEX_DISASSEMBLY |
| D | GO_CARBOHYDRATE_DERIVATIVE_BIOSYNTHETIC_PROCESS |
| D | GO_NUCLEOSIDE_MONOPHOSPHATE_METABOLIC_PROCESS |
| D | GO_ELECTRON_TRANSPORT_CHAIN |
| D | GO_OXIDOREDUCTASE_COMPLEX |
| D | GO_LIPOPROTEIN_BIOSYNTHETIC_PROCESS |
| D | GO_TRANSLATIONAL_TERMINATION |
| D | GO_MITOCHONDRIAL_MATRIX |
| D | GO_RESPIRATORY_CHAIN |
| D | GO_OXIDOREDUCTASE_ACTIVITY |
| D | GO_OXIDATION_REDUCTION_PROCESS |
| D | GO_NUCLEOSIDE_TRIPHOSPHATE_METABOLIC_PROCESS |
| D | GO_MITOCHONDRIAL_PROTEIN_COMPLEX |
| D | GO_GLUTATHIONE_METABOLIC_PROCESS |
| D | GO_MACROMOLECULAR_COMPLEX_DISASSEMBLY |
| D | GO_PEPTIDE_METABOLIC_PROCESS |
| D | GO_MITOCHONDRIAL_MEMBRANE_PART |
| D | GO_PURINE_NUCLEOSIDE_BIOSYNTHETIC_PROCESS |
| D | GO_CARBOHYDRATE_DERIVATIVE_METABOLIC_PROCESS |
| D | GO_NADH_DEHYDROGENASE_COMPLEX |
| D | GO_TRANSLATIONAL_ELONGATION |
| D | GO_MITOCHONDRION_ORGANIZATION |
| D | GO_MITOCHONDRIAL_RESPIRATORY_CHAIN_COMPLEX_I_BIOGENESIS |
| D | GO_ORGANELLAR_LARGE_RIBOSOMAL_SUBUNIT |
| D | GO_PURINE_CONTAINING_COMPOUND_METABOLIC_PROCESS |
| D | GO_CELLULAR_COMPONENT_DISASSEMBLY |
